# 4-hydroxy-2-nonenal antimicrobial toxicity is neutralized by an intracellular pathogen

**DOI:** 10.1101/2020.05.25.115097

**Authors:** Hannah Tabakh, Adelle P. McFarland, Alex J. Pollock, Rochelle C. Glover, Joshua J. Woodward

## Abstract

Pathogens encounter numerous antimicrobial responses, including the reactive oxygen species (ROS) burst. ROS-mediated oxidation of host membrane poly-unsaturated fatty acids (PUFAs) generates the toxic alpha-beta carbonyl 4-hydroxy-2-nonenal (4-HNE). Though studied extensively in the context of sterile inflammation, 4-HNE’s role during infection remains limited. Here we found that 4-HNE is generated during bacterial infection and that the intracellular pathogen *Listeria monocytogenes* induces a specific set of genes in response to 4-HNE exposure. A component of the *L. monocytogenes* 4-HNE response is the expression of the genes *rha1* and *rha2* which code for two NADPH-dependent oxidoreductases that collectively counter 4-HNE toxicity. Heterologous expression of *rha1/2* in *Bacillus subtilis* significantly increased bacterial resistance to 4-HNE both *in vitro* and following phagocytosis by murine macrophages. Our work demonstrates that 4-HNE is a previously unappreciated component of ROS-mediated toxicity and that *L. monocytogenes* has evolved specific countermeasures to survive within its presence.

## Introduction

Innate immune detection of bacterial infection initiates a complex inflammatory response characterized by production of cytokines and small molecule mediators involved in driving antimicrobial immunity. A key aspect of intrinsic cellular immunity is the production of highly reactive molecules, including reactive oxygen (ROS) and nitrogen (RNS) species (Nathan and Cunningham-Bussel 2013). Unlike the highly specific targeting of infectious agents by adaptive immune responses, ROS and RNS exhibit indiscriminate toxicity toward biological systems through their capacity to react with lipid, amino acid, and nucleic acid moieties that are conserved among both eukaryotic host and invading microbe (Patel et al. 1999, Jacobson 1996). While such indiscriminate noxious metabolite production provides protection against infection, many bacterial pathogens have evolved a diverse array of mechanisms to directly detoxify or repair damaged cellular components following ROS and RNS encounters (Fang 2004, Staerck, et al. 2017).

ROS and RNS encompass a broad group of distinct molecules. While nitric oxide, hydrogen peroxide, hypochlorite, and superoxide are well characterized molecular components of the innate immune response, these molecules give rise to numerous secondary metabolites that may also contribute to host defense against infection. An initial characteristic of the inflammatory response is the mobilization of arachidonic acid from cellular membranes. While typically involved in generation of eicosanoids, arachidonic acid exposure to oxygen radicals derived from the ROS burst leads to peroxide-mediated structural rearrangement and the generation of breakdown products, the best studied of which is 4-hydroxy-2-nonenal (4-HNE), a highly reactive membrane-permeable molecule (Hanna and Hafez, 2018). Over the last 40 years, the production of 4-HNE has been well documented at sites of sterile inflammation and has been associated with many disease pathologies, including atherosclerosis (Uchida et al., 1994), Alzheimers’ (Sayre et al., 2002), diabetes (Pillon et al., 2012), obstructive pulmonary disease (Rahman et al., 2002) and chronic liver disease (Paradis. et al., 1997). Although arachidonic acid mobilization and reactive oxygen species generation are well-known and well-studied components of innate immune responses, there are a dearth of studies characterizing the role of 4-HNE during infectious disease.

In this study, we demonstrate that 4-HNE is generated during bacterial infection both in cell culture and *in vivo*, is able to penetrate the bacterial cell envelope and access the cytoplasm, and that 4-HNE generation leads to bacterial growth delay or death. We observed that the intracellular bacterial pathogen *Listeria monocytogenes* is highly resistant to the bactericidal effects of 4-HNE and that several specific transcriptional responses are induced in response to toxic 4-HNE exposure, including two enzymes, *rha1* and *rha2*, whose loss sensitizes *L. monocytogenes* to 4-HNE toxicity. When *rha1/2* are expressed in the 4-HNE sensitive and avirulent organism *B. subtilis*, they significantly increase bacterial survival in the presence of 4-HNE both *in vitro* and following phagocytosis by murine macrophages. Our findings are consistent with the premise that 4-HNE is a heretofore unrecognized component of ROS-mediated antimicrobial defense and that pathogens have specific and multifaceted detoxification systems used to evade 4-HNE-mediated cytotoxicity in order to promote infection.

## Results

### 4-HNE accumulates during *L. monocytogenes* infection

4-HNE is a highly reactive electrophilic αβ-unsaturated aldehyde that undergoes Michael addition with nucleophilic amino acids, resulting in stable conjugates that correlate with cellular levels of free 4-HNE. Monoclonal antibodies to these adducts are routinely used to monitor 4-HNE levels in cells (Majima, Nakanishi-Ueda, and Ozawa. 2002). To investigate 4-HNE production during bacterial infection, we infected murine hepatocytes with *L. monocytogenes* and quantified 4-HNE conjugates using dot blots of whole cell lysates at various times post-infection. During the course of a 24-hour infection, a continuous increase in 4-HNE protein conjugates relative to β-actin control was observed (Fig 1A). To interrogate the impact of bacterial infection on host production of 4-HNE *in vivo*, mice were infected intravenously with *L. monocytogenes* constitutively expressing GFP. At 48 hours post infection, tissues were harvested, fixed, and analyzed by immunohistochemistry. Clear foci of infection were visible in the liver with no change in the abundance of 4-HNE protein conjugates (Supp Fig 1A-D). In the spleen, however, the bacteria were diffusely distributed throughout the organ and the entire spleen of the infected mouse exhibited increased staining for 4-HNE protein conjugates (Figs 1B-E).

**Fig 1.**
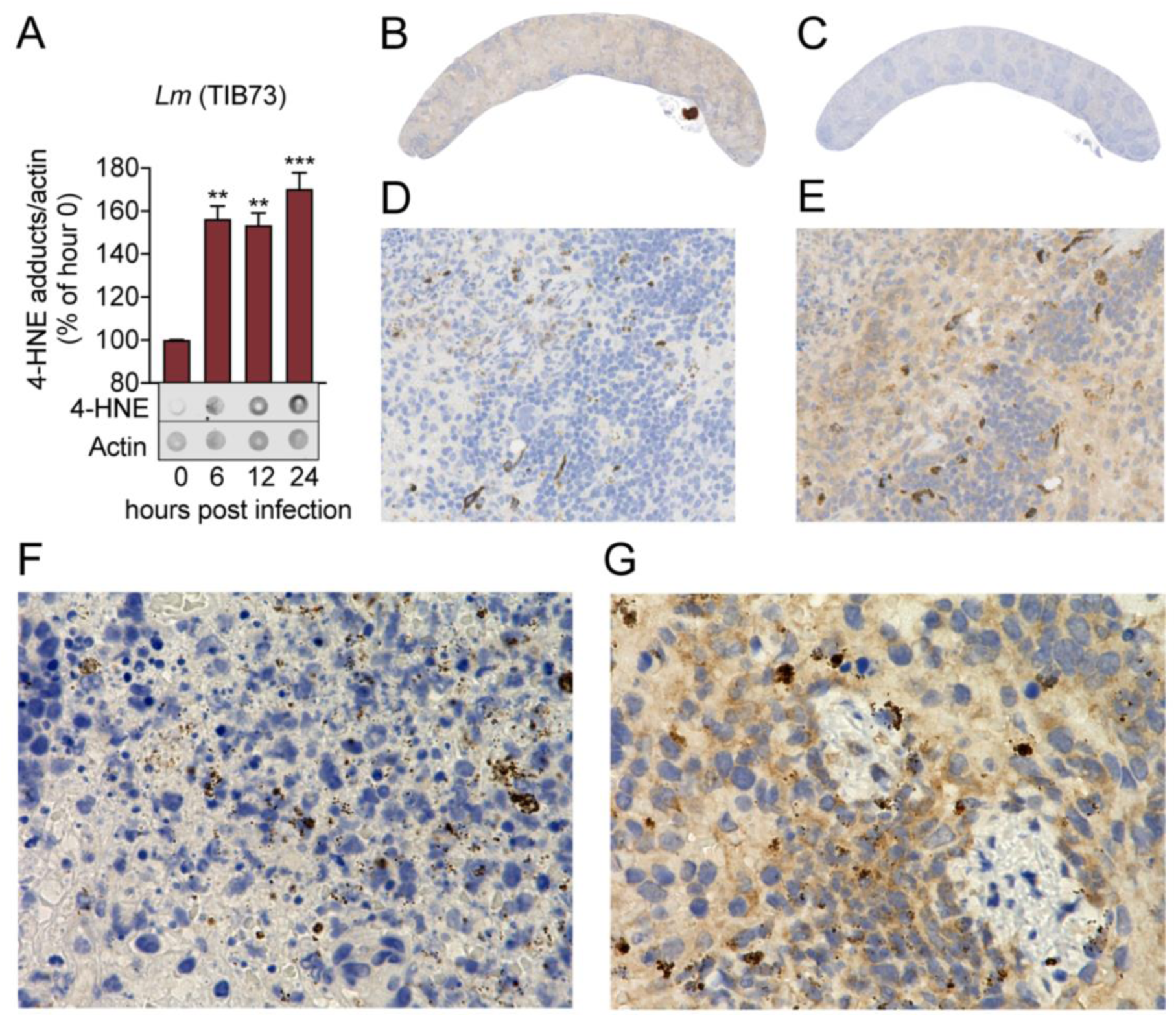
4-HNE accumulates during intracellular bacterial infection by *L. monocytogenes*. (A) 4-HNE accumulation in TIB73 murine hepatocytes during intracellular *L. monocytogenes* infection. 4-HNE adduct accumulation is assessed by dot blot whole cell Western and normalized to actin levels. Data are normalized 4-HNE levels as percent of 4-HNE at hour 0 of infection. Western image below is representative. (B) 4-HNE accumulation during 48-hour murine infection by GFP_+_ *L. monocytogenes* assessed by immunohistochemistry analysis. Spleen (infected) anti-4-HNE. (C) spleen (uninfected) anti-4-HNE. (D) spleen (infected) anti-GFP 25x magnification. (E) spleen (infected) anti-4-HNE 25x magnification (F) spleen (infected) anti-GFP 100x magnification. (G) spleen (infected) anti-4-HNE 100x magnification. Antigens detected with 3,3-diaminobenzidine staining by horseradish peroxidase and cellular nuclei imaged with Hematoxylin counterstain. Data in (A) are pooled from two independent experiments. Statistics in (A) are an ordinary one-way ANOVA with a Dunnett’s multiple comparison test against hour 0. Error bars are mean +/- SD. *, p < 0.05; **, p < 0.01; ***, p < 0.001.

The grossly different observations between the liver and spleen were somewhat surprising given our previous observation that infection of murine hepatocytes with *L. monocytogenes in vitro* induced accumulation of 4-HNE conjugates. These observations are likely a consequence of several factors. The liver is a major site of small molecule detoxification and hepatocytes are known to produce high levels of many of the 4-HNE metabolizing proteins, including aldo-keto reductases, alcohol dehydrogenase, and the 4-HNE glutathione transferase, GSTA4 (Zheng et al, 2014). Additionally, the immune driven mobilization of arachidonic acid and ROS precursors that lead to 4-HNE generation may result in elevated accumulation of this host aldehyde in the spleen.

At higher levels of magnification, we observed that 4-HNE conjugates were not evenly distributed among all cells. The majority of cells exhibited diffuse and constant staining and a subset showed very dark and robust staining for 4-HNE conjugates. At 100x magnification, the most pronounced signal for 4-HNE conjugates had punctate staining resembling the staining of the bacteria with anti-GFP antibody (Fig 1F-G), consistent with 4-HNE conjugate formation on the bacteria. While these observations do not provide quantitative measures of 4-HNE levels, they establish that 4-HNE was indeed elevated following bacterial infection and suggest that bacteria encounter and may be directly chemically modified by this metabolite during infection.

### 4-HNE causes damage through the targeting of nucleophilic protein moieties and *L. monocytogenes* is uniquely resistant to 4-HNE-mediated death

Electrophilic stress due to 4-HNE conjugation to proteins causes eukaryotic cells to undergo apoptosis following intermediate 4-HNE exposure (5-40µM) and necrosis at higher concentrations (40-100µM) (Dalleau et al., 2013). However, due to 4-HNE’s lipophilicity it is believed to accumulate to significantly higher levels (0.3-5mM) near and within membranes than what is typically considered cytotoxic (Zimniak, 2011; Esterbauer, Schaur and Zollner, 1991; Uchida, 2003).

To characterize bacterial sensitivity to 4-HNE toxicity, we exposed bacteria within the Firmicutes phylum to a wide range of 4-HNE concentrations and assessed viability. We observed variability in survival, ranging from a 4-log reduction in CFU for *B. subtilis*, a 2-log reduction for *Staphylococcus aureus*, a half-log reduction for *Enterococcus faecalis* to less than a 40% reduction for *L. monocytogenes* (Fig 2A). We observed a dose dependent decrease in growth of *L. monocytogenes* with increased exposure to 4-HNE (Fig 2B). The significant resistance of *L. monocytogenes* to 4-HNE exposure was unexpected. Due to the conserved nature of 4-HNE targets, we hypothesized that 4-HNE would exert similar damaging effects on bacteria as on eukaryotic cells. Thus, we first interrogated the ability of 4-HNE to generate protein adducts in *L. monocytogenes*. Dot blots of *L. monocytogenes* bacterial cell lysates indicated an increase in 4-HNE-protein adducts that correlated with increased 4-HNE exposure (Fig 2C), establishing that this aldehyde penetrates the bacterial cell envelope and impacts cytosolic proteins.

**Fig 2.**
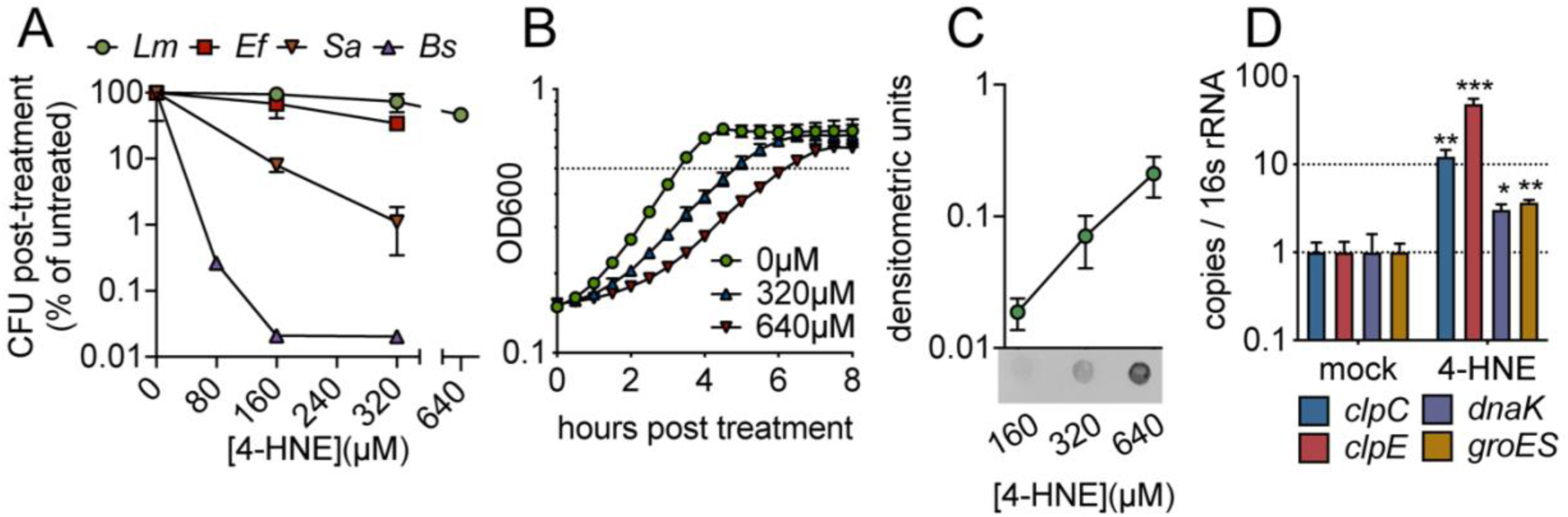
4-HNE impacts bacterial viability, replication and damage to cellular proteins. (A) Survival of mid-log (0.4-0.8OD) *Listeria monocytogenes (Lm), Enterococcus faecalis (Ef), Staphylococcus aureus (Sa) and Bacillus subtilis (Bs)* following exposure to various concentrations of 4-HNE or mock vehicle (ethanol) in PBS at 37 °C for 1 hour. Data are reported as recovered CFU normalized to mock treated controls. (B) Growth of *L. monocytogenes* in TSB at 37°C with various concentrations of 4-HNE added at time zero. (C) Anti-4-HNE dot blot of soluble bacterial lysates from mid-log *L. monocytogenes* suspended in PBS and treated with increasing concentrations of 4-HNE for 30 minutes. Protein levels were normalized (3µg total protein) and signal quantified by densitometry on a Licor Odyssey Fc. (D) qRT-PCR measurement of expression of indicated genes in midlog *L. monocytogenes* in TSB treated with 640 µM 4-HNE for 20 minutes. Expression normalized to 16S rRNA levels. Data in figures (A) and (B) are in biological triplicate. Data in (C) and (D) are biological duplicate. Statistics for (D) are unpaired t-tests of ΔΔCt values of treated samples versus untreated samples for each gene. Error bars are mean +/- SD. *, p < 0.05, p < 0.01; ***, p < 0.001.

4-HNE adduct accumulation can result in protein misfolding and crosslink-induced aggregation. Eukaryotic cells clear 4-HNE damaged proteins through proteasome and autophagy-mediated pathways (Zhang and Forman, 2017). Bacteria target damaged proteins for degradation through the proteases that comprise the heat shock response (Parsell and Lindquist, 1993). This pathway is primarily transcriptionally regulated (Yura, Nagai and Mori, 1993), so RT-qPCR was performed on a subset of heat shock genes representing two major groups of heat shock genes *in L. monocytogenes*: HrcA-regulated chaperones and CtsR-regulated proteases (Roncarati and Scarlato, 2017). When *L. monocytogenes* was exposed to 640µM 4-HNE for 20 minutes, the four genes tested (*clpC, clpE, dnaK, groES*) were significantly induced compared to vehicle controls (Fig 2D). These data, combined with the dot blot results, support the hypothesis that 4-HNE causes protein damage to which *L. monocytogenes* mounts a heat shock response. The elevated induction of cellular proteases relative to chaperones may indicate that *L. monocytogenes* primarily combats electrophilic 4-HNE stress through turnover of damaged proteins rather than chaperone-mediated stabilization. Collectively, these observations suggest that despite formation of protein adducts and delayed growth following exposure, *L. monocytogenes* has a uniquely robust capacity to survive 4-HNE toxicity even among closely related organisms.

### *L. monocytogenes* expresses potential 4-HNE detoxification enzymes that contribute to its survival in the presence of 4-HNE

Our data suggest that *L. monocytogenes* is exposed to 4-HNE during infection and that only high concentrations of this aldehyde impact its growth. We hypothesized that *L. monocytogenes* may express genes involved in countering the cytotoxic effects of 4-HNE. To probe further, we performed global transcriptome analysis during 4-HNE exposure using RNA sequencing. Over one hundred genes were induced greater than 10-fold in response to 4-HNE exposure, including several of the heat shock genes previously identified by qRT-PCR analysis (Fig 3A, Suppl Fig 2A, Suppl Table 1).

**Fig 3.**
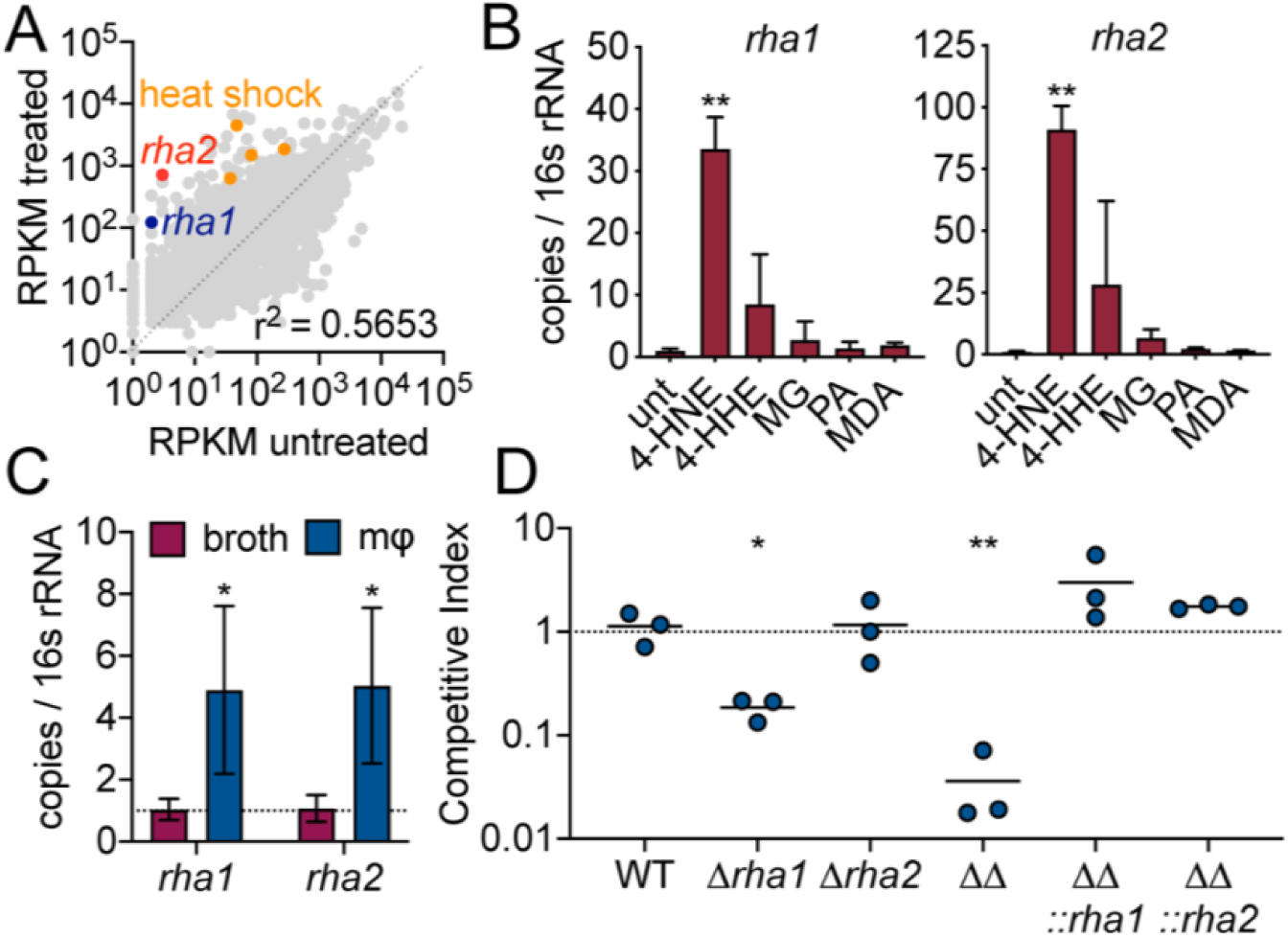
4-HNE exposure induces resistance genes in *L. monocytogenes.* (A) Global gene expression of midlog *L. monocytogenes* in TSB treated with 640 µM 4-HNE or ethanol control for 20 minutes. RPKM: reads per kilobase million. Genes of interest *rha1, rha2* and heat shock class members are indicated in blue, red and orange, respectively. (B) qRT-PCR of expression of *rha1* and *rha2* genes after 20-minute treatment of mid-log bacteria in TSB media with 500µM of selected aldehydes: 4-HNE (4-hydroxy-2-nonenal), 4-HHE (4-hydroxy-2-hexenal), MG (methylglyoxal), PA (propionaldehyde) and MDA (malondialdehyde). (C) qRT-PCR analysis of expression of *rha1* and *rha2* at 6 hours post infection in J774 macrophages (mφ). (D) Competitive index of midlog WT and mutant *L. monocytogenes* in PBS treated with 640 µM 4-HNE at 37°C for one hour. Data in figures (A) and (B) are in biological duplicate. (C) is in biological triplicate. (D) is in technical triplicate, representative of at least two independent experiments. Statistics in (B) is an ordinary one-way ANOVA with Dunnett’s multiple comparison test of ΔΔCt values of the treated samples against untreated samples. Statistics in (C) are unpaired t-test between broth and macrophage samples. Statistics in (D) are unpaired t-test between WT and mutant *L. monocytogenes* competition pairs. Error bars are mean +/- SD. *, p < 0.05; **, p <0.01.

Eukaryotic cells utilize several reductases to detoxify 4-HNE (Mol et al., 2017). Two reductases were highly induced in our global transcriptome, *lmo0103* and *lmo0613*, which we refer to as *rha1* and *rha2* (**r**esistance to **h**ost **a**lkenals 1 and 2), respectively. Phyre2 analysis of Rha1 predicted high structural homology to CLA-ER, a flavin-dependent enone reductase from *Lactococcus plantarum* (PDB: 4QLY)(Hou et al, 2015). A Phyre2 analysis of Rha2 revealed structural similarity to a few crotonyl-CoA carboxylase/reductases (PDBs: 3KRT, 4Y0K, and 5A3J) and a plant chloroplast oxoene reductase (PDB: 5A3V), two enzymes with the capacity to reduce enone-containing lipophilic substrates. Given the predicted reductase activity of Rha1 and Rha2 and their structural similarity to proteins that metabolize enone-containing compounds, we further investigated their role in 4-HNE resistance.

Induction of *rha1* and *rha2* in response to 4-HNE exposure was found to be 34 and 90-fold, respectively, by RT-qPCR. To assess the specificity of their induction, we exposed *L. monocytogenes* to a sublethal concentration of a panel of aldehydes (Suppl Fig 2B,C), including 4-HNE; 4-HHE (4-hydroxy-2-hexenal), a similar but shorter chain αβ-unsaturated aldehyde produced from the oxidation of ω-3 fatty acids (Awada et al., 2012); methylglyoxal, a reactive byproduct of glycolysis; propionaldehyde, an alpha hydrogen aldehyde; and malondialdehyde, another product of lipid peroxidation (Esterbauer, Schaur, and Zollner, 1991). Both *rha1* and *rha2* were most strongly induced by 4-HNE exposure, with much less induction by 4-HHE and negligible induction with the other tested compounds (Fig 3B). We next assessed if these genes are induced by *L. monocytogenes* during intracellular infection. At 6 hours post-infection of macrophages we found that there was significant induction of both genes compared to growth in BHI broth (Fig 3C). Together these transcriptional studies suggested that the *rha1/2* genes are not components of a general aldehyde response but rather specific responses to 4-HNE encountered during infection.

These intriguing transcriptional results suggested that *rha1* and *rha2* may function in 4-HNE resistance. Individual and double mutants of these genes were generated and assessed by competition experiments for survival relative to WT *L. monocytogenes* following 4-HNE exposure. Control mixtures left untreated in PBS exhibited no significant difference between mutant and control strains (Suppl Fig 2D).

Among mixtures exposed to 640 µM 4-HNE, unmarked WT and marked WT showed no significant difference in 4-HNE survival. Loss of *rha2* had no effect on 4-HNE survival while Δ*rha1* had a modest 5-fold reduction relative to WT. However, the *Δrha1Δrha2* mutant exhibited a 50-fold competitive defect compared to WT *L. monocytogenes* that was rescued by either *rha1* or *rha2* expression in *trans*, demonstrating that both genes must be absent for the toxic effect to manifest (Fig 3D). The *Δrha1Δrha2 L. monocytogenes* was used to infect macrophages and mice. During cellular infection, we observed a ∼50% reduction in CFU at 2 hours post-infection (Suppl fig 2E), which rebounded to WT levels at later time points. In mice infected via intravenous injection no significant phenotype was observed at 48 hours post infection in either the spleen or the liver (Suppl fig 2F). These findings provide evidence that Rha1 and Rha2 are specific and important for 4-HNE resistance, though other factors likely contribute to *L. monocytogenes* capacity to counteract this metabolite *in vivo*.

### Recombinant Rha1 and Rha2 metabolize 4-HNE and ectopic expression confers 4-HNE resistance to sensitive bacteria

In order to determine if these putative enone reductases can directly utilize 4-HNE as a substrate, we generated recombinant Rha1 and Rha2 proteins. As controls for these studies, we generated catalytically dead variants of the two proteins by mutating amino acids predicted to be involved in flavin or NADPH binding (Suppl Fig 3A). All proteins were expressed and characterized for NADPH oxidation in the presence and absence of 4-HNE. Only the WT variants of Rha1 and Rha2 exhibited NADPH oxidation upon addition of 4-HNE (Fig 4A), consistent with their capacity to mediate NADPH dependent reduction of the enone.

**Fig 4.**
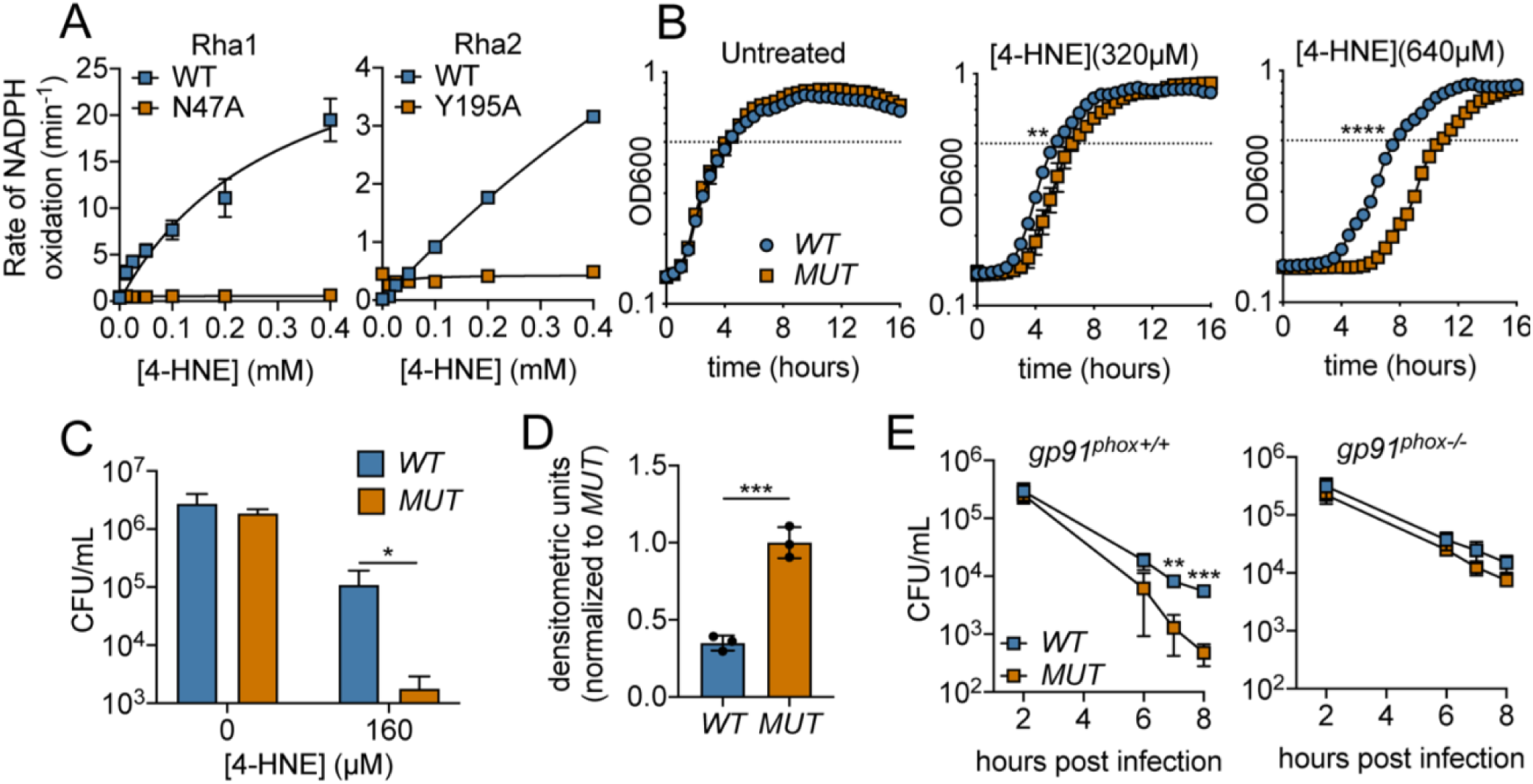
Recombinant Rha1 and Rha2 metabolize 4-HNE and ectopic expression confers 4-HNE resistance to sensitive bacteria. (A) Rates of NADPH oxidation (200µM) by WT and mutant variants of Rha1 and Rha2 in the presence of 4-HNE. (B) Growth of *B. subtilis* expressing either the *WT* or *MUT* (catalytically dead mutant) versions of *rha1* and *rha2* in TSB at 37 °C in the presence of the indicated concentrations of 4-HNE added at time zero. Dotted line represents OD_600_ 0.5. (C) Survival of *B. subtilis WT* and *MUT* in PBS at 37 °C with 160µM 4-HNE for one hour. (D) 4-HNE conjugates from *B. subtilis WT* and *MUT* soluble cell lysates (3 µg total protein) three hours after 4-HNE treatment as assessed by dot blot and quantified by densitometry on a Licor Odyssey Fc. (E) *B. subtilis WT* and *MUT* survival following phagocytosis by Interferon gamma-activated primary WT or phagosomal oxidase-deficient bone marrow derived macrophages (WT or *gp91*^*phox-/-*^ pBMMs). All experiments in figure performed in biological triplicate. Statistics in (A) are unpaired t-tests comparing the hours to OD 0.5 between *WT* and *MUT B. subtilis pHT01::rha1/2*. Statistics in (B), (C) and (D) are unpaired t-tests comparing *WT* and *MUT B. subtilis pHT01::rha1/2*. Error bars are mean +/- SD. *, p < 0.05; **, p < 0.01; ***, p < 0.001; ****, p < 0.0001.

We hypothesized that Rha1 and Rha2 could confer 4-HNE resistance to a sensitive organism. To this end, each gene and corresponding catalytically dead variant were expressed individually and in combination in *B. subtilis*, which exhibited 300-fold more sensitivity than *L. monocytogenes* upon 4-HNE exposure (Fig 2A, suppl fig 3B). Consistent with our observations of the *L. monocytogenes rha* deletion mutants, expression of *rha2* in *B. subtilis* had no effect on growth in the presence of 4-HNE, *rha1* gave modest protection against 640µM 4-HNE, and expression of both *rha1* and *rha2* had the largest growth rescue, reducing lag time by up to 3 hours (Suppl Fig 3C, Fig 4B). We then focused on *B. subtilis* expressing both *rha1* and *rha2* genes, as this strain had the most robust phenotype. When assessed for bacterial survival following 4-HNE treatment, *B. subtilis* expressing the functional enzymes exhibited nearly a 2-log survival advantage relative to the control strain expressing enzymatically dead *rha1/2* (Fig 4C). Additionally, soluble cellular fractions from *B. subtilis* exposed to 4-HNE and probed for 4-HNE protein adducts by dot blot revealed a ∼70% reduction in 4-HNE conjugates in the *B. subtilis* strain expressing both of the active *rha1* and *rha2* genes versus their catalytically dead counterparts (Suppl Fig 3D, Fig 4D).

To determine if 4-HNE resistance conferred by *rha1/2* could contribute to bacterial survival within mammalian cells, *B. subtilis rha1/2* strains were assessed for viability following phagocytosis by primary bone marrow-derived macrophages. We found that *B. subtilis* expressing the active forms of Rha1 and Rha2 maintained a significantly higher CFU over the course of eight hours than the *B. subtilis* expressing the catalytically dead form (Fig 4E). To determine whether this survival advantage was due to 4-HNE resistance, we measured *B. subtilis* survival within bone marrow-derived macrophages from *gp91*^*phox-/-*^ mice deficient in oxidase cytochrome b-245, which are unable to produce the reactive oxygen burst and therefore 4-HNE (Esterbauer, Schaur, and Zollner, 1991). Consistent with the role of *rha1* and *rha2* in resistance to a ROS-derived factor, the protective effect of WT *rha1/2* expression was eliminated in the absence of *gp91*^*phox*^ (Fig 4E). Together, these observations revealed that expression of *rha1* and *rha2* in *B. subtilis* imparts resistance to 4-HNE toxicity and impacts bacterial survival in the host cell’s ROS burst.

## Discussion

In this study, we provide evidence that ROS-derived metabolite 4-HNE accumulates during *L. monocytogenes* infection both in tissue culture and in mice. We also show that 4-HNE exhibits antimicrobial effects on several bacterial species and that in the highly resistant intracellular pathogen *L. monocytogenes*, exposure to this aldehyde induces a specific and robust transcriptional profile. Among the highest induced genes are components of the heat shock response, consistent with aldehyde induced protein damage, a known effect of 4-HNE exposure. In addition, two genes, *rha1* and *rha2* are highly and specifically induced by 4-HNE exposure, and these two enzymes turnover 4-HNE through NADPH-dependent reduction *in vitro*. Disruption of *rha1* and *rha2* in *L. monocytogenes* results in a decrease in viability in the presence of 4-HNE, and heterologous expression of *rha1* and *rha2* in *B. subtilis* conferred increased tolerance to 4-HNE toxicity in this non-pathogenic and 4-HNE-sensitive organism. Rha1 and Rha2 expression in *B. subtilis* also allowed for greater survival following phagocytosis by bone marrow derived macrophages in a manner entirely dependent upon phagocyte ROS generation. Together this work supports the conclusion that 4-HNE represents one of the individual molecular components of ROS-mediated host defense through its direct antimicrobial effects on bacteria and that pathogens have likely evolved complex mechanisms of surviving its encounter within eukaryotic hosts (Fig 5).

**Figure 5.**
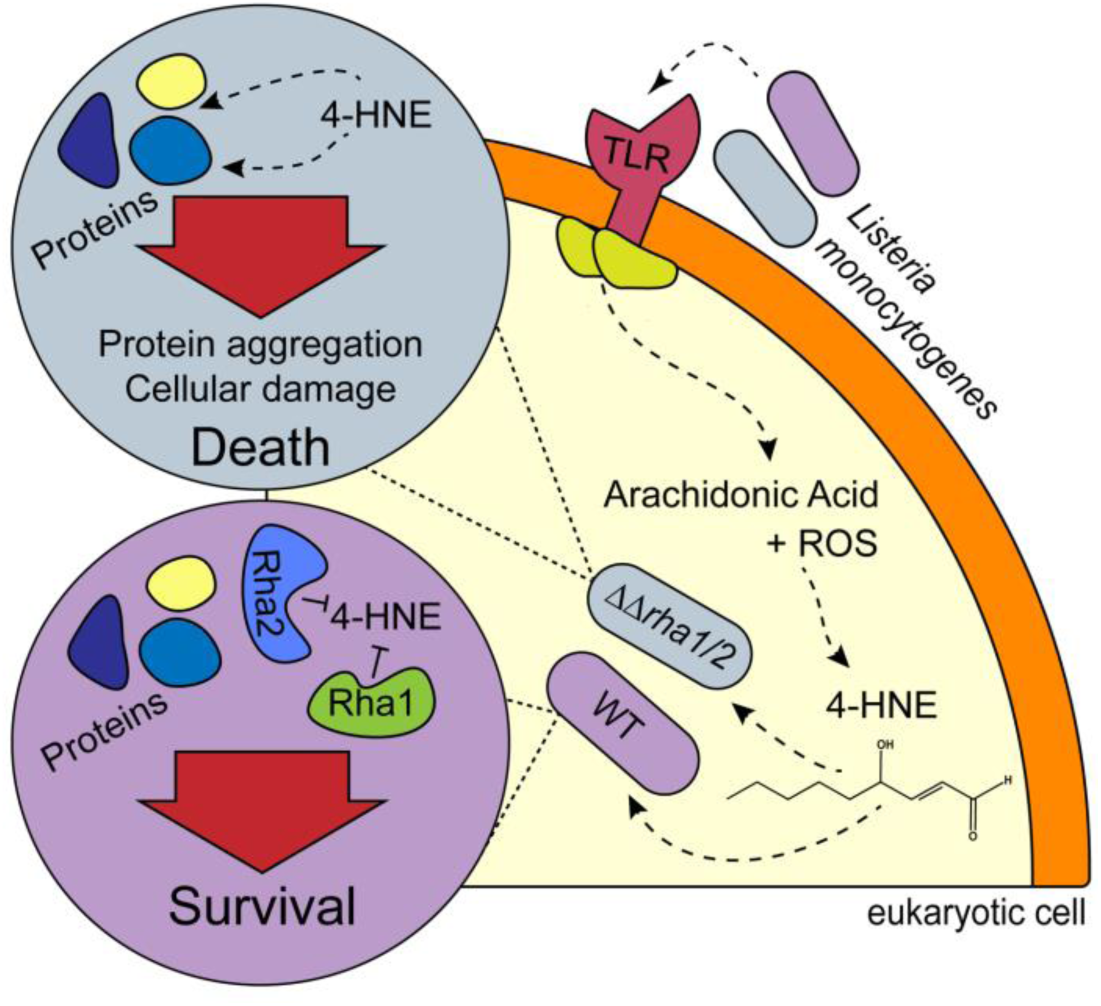
Model of *rha1/2*-mediated 4-HNE resistance in *L. monocytogenes.*

There are many parallels between the chemical and biological functions of 4-HNE and other toxic metabolites that function in antimicrobial defense. The freely diffusible and highly reactive diatomic gas nitric oxide (NO) is produced during infection (Iyengar, Stuehr and Marletta, 1987; Stuehr and Marletta, 1985) and has a direct role in preventing bacterial growth (Nathan and Hibbs, 1991). However, due to the conservation of its reactive targets, elevated levels of NO also exert pathological effects during both sterile inflammation and acute infections (Nagafuji et al., 1995, Galley and Webster, 1998). 4-HNE is membrane diffusible, highly reactive, and contributes to disease pathology due to its cytotoxic activity toward eukaryotic cells. These parallels, together with our findings that bacterial infection induces 4-HNE production are consistent with the premise that 4-HNE represents a component of ROS-mediated host defense, among such other toxic metabolites as superoxide, hydrogen peroxide, and hypochlorite.

While a role for 4-HNE in host antimicrobial defense has yet to be appreciated in mammals, plants utilize a variety of lipophilic molecules generated by the oxidation of polyunsaturated fatty acids (PUFAs), collectively referred to as oxylipins. While generally considered to be involved in signal transduction, many oxylipins can directly inhibit bacterial growth (Prost et al., 2005) and 4-HNE itself is a component of the oxylipin burst in soybean where it serves an anti-fungal function (Vaughn and Gardner, 1993). Because 4-HNE is one of several distinct metabolites produced following oxidation of PUFAs in mammals, it is conceivable that other reactive byproducts of this process also contribute to microbial defense in a similar manner.

To survive within the sterile tissues of eukaryotic hosts, bacterial pathogens often counteract the toxic effects of the immune response. Our discovery of two genes that confer synergistic resistance to 4-HNE in *L. monocytogenes* begin to provide insight into the mechanisms by which 4-HNE toxicity might be overcome. *In vitro* studies suggest that Rha1 and Rha2 both metabolize 4-HNE in an NADPH-dependent manner, suggesting redundant functions. Redundancy in bacterial resistance to ROS is a relatively common phenomenon, including the need to eliminate five individual enzymes in *Salmonella enterica* Serovar Typhimurium to exhibit a phenotype in the presence of hydrogen peroxide (Hébrard et al., 2009) and simultaneous disruption of four enzymes in *Bacillus anthracis* to observe a phenotype in the presence of superoxide (Cybulski et al., 2009). Such redundancy in bacterial detoxification programs likely decreases the chances that genetic drift or other genomic damage would render an organism defenseless against oxidative stress.

While Rha1 and Rha2 both contribute to 4-HNE resistance, the *Δrha1Δrha2 L. monocytogenes* strain still exhibits several logs of survival benefit relative to the related organism *B. subtilis*, suggesting that other mechanisms of 4-HNE resistance remain to be identified. Among the many uncharacterized genes induced during 4-HNE exposure, *lmo0796* shows homology to *bcnA*, a secreted lipocalin in *Burkholderia cenocepacia* which sequesters long-chain lipophilic antibiotics (El-Halfawy et al., 2017). 4-HNE, with its long hydrophobic tail, could conceivably be neutralized in an analogous manner. It is possible many intrinsic resistance properties of *L. monocytogenes* are not reflected through transcriptional responses. For instance, addition of amine containing constituents on the cell’s surface through lysinylation of teichoic acids and/or lipids, as well as deacetylation of peptidoglycan, may provide a nucleophile reactivity barrier that prevents 4-HNE entry into the bacterial cell. Additionally, αβ-unsaturated aldehydes have preferential reactivity toward sulfhydryl groups, including cysteine and glutathione, and it is expected that thiolate depletion would be the major mechanism of 4-HNE toxicity (Lopachin and Gavin, 2014). Indeed, the thiol responsive transcription factor spxA1 was induced >2-fold in response to 4-HNE and the magnitude of heat shock gene induction mirrored results reported for *B. subtilis* following diamide treatment, a potent inducer of disulfide stress (Ole Leichert, Scharf, and Hecker, 2003). While our findings provide initial molecular insight into one pathogen’s resistance to 4-HNE, it is clear that many details are yet to be revealed.

Taken together, our findings extend the range of antimicrobial molecules generated through the reactive oxygen burst to include the byproducts of lipid peroxidation. Additionally, bacteria whose infection cycles involve intimate exposure to these molecules, such as *L. monocytogenes*, have specific and finely regulated detoxification and protection programs against this toxicity. Future investigation of the impacts of 4-HNE on a diverse array of organisms with varied infection models will highlight the importance of this metabolite on host defense and the varied mechanisms by which pathogens counteract its toxicity to promote infection.

## Acknowledgements

We thank Aruna Menon, Maureen Thomason and Qing Tang for assistance with experiments, Brian Johnson for performing the histology experiments at the Histology and Imaging Core at the University of Washington, Steven Libby for providing gp91 knockout mice and Peter Lauer and Kevin Lang for providing plasmids. We thank members of the Woodward-Reniere supergroup for helpful discussions, and the Mougous, Reniere and Lagunoff labs for reagents. This material is based upon work supported by PHS NRSA T32GM007270 from NIGMS (to HT and AJP), National Science Foundation Graduate Research Fellowship Program under grant no. DGE-1256082 (to APM), and National Institutes of Allergy and Infectious Disease grants R01AI116669 and R21AI127833 (to JJW).

## Author Contributions

HT, APM and JJW conceptualized the studies, HT, APM, AJP and RCG performed the experiments, HT and JJW wrote the original draft of the manuscript, APM, AJP and RCG edited and reviewed the manuscript, and JJW, HT, AJP and APM acquired funding. JJW supervised and administered the project.

## Methods

### Strains and routine growth conditions

Unless otherwise specified, *L. monocytogenes* was grown at 37°C degrees in tryptic soy broth (TSB) media and E. coli at 37°C degrees in Luria-Bertani (LB) media with appropriate antibiotic selection. Unless otherwise noted, *B. subtilis* was struck on LB plates with appropriate antibiotics and induction agent overnight at 30°C, after which the biomass was scraped off the plates, resuspended in LB media, passed 6 to 10 times through a 27-gauge needle to break up clumps and chains, then normalized to an OD_600_ of 1.

When required for selection, antibiotic concentrations used in this study were as follows – *L. monocytogenes* selections: streptomycin 200µg/mL, chloramphenicol 5µg/mL; *E. coli* selections: ampicillin 50µg/mL; *B. subtilis* selections: chloramphenicol 10µg/mL; tissue culture: gentamicin 50µg/mL.

### DNA manipulation and plasmid construction

All DNA manipulation procedures followed standard molecular biology protocols. Primers were synthesized and purified by Integrated DNA Technologies (IDT). HiFi polymerase (Kapa Biosystems, #KK2102), FastDigest restriction enzymes (Thermo Fisher Scientific #FD0274) and T4 DNA ligase (Thermo Scientific # K1423) were used for plasmid construction, with the exception of *pHT01::rha1/2_WT* and *pHT01::rha1/2_MUT* which were generated using Gibson Assembly MasterMix (NEB, #E2611S). DNA sequencing was performed by Genewiz Incorporated.

### Bacterial infection dot blot

TIB73 cells were lysed in whole cell lysis buffer (50 mM Tris pH 7.5, 150 mM NaCl, 1% Triton X-100) with EDTA (1 µM) and Halt Protease Inhibitor Cocktail (Thermo Fisher Scientific, #78442). Protein concentration was determined using the Pierce BCA protein assay kit (Fisher Scientific, #PI23227). Lysates were resuspended in 1X PBS to achieve 2µg of protein per 3µl, which was the volume spotted out onto nitrocellulose membrane (Bio-Rad, #1620115). Four to six technical replicates were spotted per biological replicate. Membranes were blocked in 1X TBS-T (Tris-buffered saline with 0.1% Triton X-100) with 5% BSA (sterile filtered) for 1 hour at room temperature, incubated for 1 hour with anti-4-hydroxynonenal (HNE) antibody (Abcam #ab46545, diluted 1:1000) and anti-actin antibody (Santa Cruz Biotechnology #sc-8432, diluted 1:500), washed and then incubated with the appropriate IRDye secondary antibodies (LI-COR). Membrane fluorescence was visualized and densitometric analyses were performed using an Odyssey CLx imager and Image Studio software (LI-COR).

### Mouse infections

*L. monocytogenes* was grown overnight statically at 30°C degrees in Brain Heart Infusion (BHI) broth, then back-diluted using 1.2 mL of overnight culture to 4.8 mL of fresh BHI and grown for 1 hour at 37°C degrees shaking. OD_600_ of these cultures were taken and, using the conversion of 1 OD = 1.7×10^9^ CFU, diluted to 5×10^5^ CFU/mL with PBS. 200µl were then injected into female mice between 6-8 weeks of age retro-orbitally (1×10^5^ CFU/mouse) and livers and spleens were harvested at 48 hours post infection. Livers were homogenized in 10mL of cold 0.1% IGEPAL and spleens were homogenized in 5mL using a Tissue Tearor Model 985370 (Biospec Products) at 10,000 RPM for 5 seconds/organ. Homogenates were diluted in PBS and plated on LB plates to enumerate CFU. All protocols were reviewed and approved by the Institutional Animal Care and Use Committee at the University of Washington.

### 4-HNE histology

Two female mice were infected as outlined above, in addition to one uninfected control mouse. The livers and spleens were harvested at 48 hours post infection and placed in 10% neutral buffered formalin for 24 hours, after which the organs were removed from formalin and placed in PBS for 24 hours. Paraffin embedded tissues were sliced and prepared as slides. Slides were then deparaffinized for 30 minutes at 60°C. All subsequent manipulations were performed on a Leica Bond Automated Immunostainer. Antigen retrieval for GFP was performed by HIER 2 (EDTA) treatment for 20 minutes at 100°C. Antigen retrieval for 4-HNE was performed by citrate treatment for 20 minutes at 100°C. Then a Leica Bond peroxide block was performed for 5 minutes at room temperature, and normal goat serum (10% in TBS) was added for 20 minutes at room temperature. Primary antibody was added (GFP 1:500), (Rab IgG 1:1000), (4-HNE 1:200) in Leica Primary antibody diluent, for 30 minutes at room temperature. Leica Bond Polymer was added for 8 min at room temperature, after which the samples were washed with Leica Bond Mixed Refine (DAB) detection solution twice for 10 minutes at room temperature. Hematoxylin Counterstain was added for 4 minutes and the samples were cleared to xylene. Finally, samples were mounted with synthetic resin mounting medium on a 1.5cm coverslip and imaged with a Hamamatsu Nanozoomer Whole Slide Scanner and a Keyence BZ-X710 Microscope.

### *L. monocytogenes* 4-HNE dot blots

*L. monocytogenes* were sub-cultured from overnight stationary cultures 1:100 into fresh media and grown to mid-log (0.4-0.8 OD_600_). The bacteria were normalized to OD 1, washed twice with sterile PBS and resuspended in sterile PBS. A range of 4-HNE concentrations were added to the bacteria and the samples were placed at 37C for 30 minutes. Upon completion, the bacteria were washed twice with PBS and spun at 10,000g for 5 minutes, then resuspended in fresh PBS. The bacteria were then sonicated using a narrow tip sonicator at 20% power, 1 second on 1 second off for 10 seconds and placed on ice. The bacteria were then spun at 4C at 10,000g for 30 minutes. The subsequent lysate was transferred to fresh Eppendorf tubes containing Halt Proteinase and Phosphatase Inhibitor (Thermo Fisher Scientific, #78442) and stored at -80C until use. For the dot blots, the protein concentration was normalized using BCA (Fisher Scientific, #PI23227) and 3µg in 3µL was spotted onto nitrocellulose membrane (Bio-Rad, #1620115). The nitrocellulose was then dried, blocked for 45 minutes in 5% dry milk, washed 3 times with TBS-T and primary 4-HNE antibody was added at 1:200 dilution (Abcam, #ab46545). The antibody was left on overnight at 4°C rocking. The primary antibody was then washed off with TBS-T 3 times and secondary antibody (Licor, #926-32211) was added at 1:8000 for 45 minutes at RT. The secondary antibody was then washed off with TBS-T twice, TBS once and the blot was imaged on a Licor Odyssey Fc. Relative densitometric analysis was performed using Licor Image Studio software.

### RNA extraction from broth cultures of *L. monocytogenes*

*L. monocytogenes* stationary phase overnights were subcultured 1:100 into fresh TSB media and grown shaking at 37°C to mid-log (0.4-0.8 OD_600_). Then a final concentration of 640µM 4-HNE (Cayman #32100) or vehicle (100% ethanol) was added to the bacteria, which continued to grow at 37°C shaking for 20 minutes. After 20 minutes, ice cold 100% methanol was added in equal volume to the culture flask and placed at -20C overnight. The next day the bacteria were spun down and resuspended in 400µL AE buffer (50mM NaOAc pH 5.2, 10mM EDTA in molecular grade water). The resuspended bacteria were then mixed with 400µL acidified 1:1 phenol:chloroform pH 5.2 (Fisher Scientific, # BP1753I) and 40µL 10% sodium dodecyl sulfate (SDS) and was vortexed for 10 minutes in a multi-tube vortexer. The tubes were then transferred to a 65C heat block for ten minutes, after which the mixture was transferred to a Heavy Phase-lock tube (VWR #10847-802) and spun down for 5 minutes at 17,000g. Then the aqueous layer was transferred into tubes containing 1mL 100% ethanol and 40µL 3M NaOAc and placed at -20C for 6 hours. Then tubes were spun at 17,000g for 30 minutes at 4C, the ethanol was aspirated and 500µL 70% ethanol was added. The tubes were then centrifuged at 17,000g for 10 minutes at room temperature and the supernatant was aspirated. The RNA pellet was then dried in a speed vacuum concentrator for 5 minutes and resuspended in RNA-free molecular grade water. The extracted RNA was then treated with DNase (Ambion Life Technologies #AM1907) for an hour at 37°C and used for downstream processing.

### RNA-sequencing

RNA was processed using the Ovation Complete Prokaryotic RNA-Seq Library System (NuGEN, #0363-32, 0326-32, 0327-32) according to the manufacturer’s instructions to a final pooled library concentration of 3nM. Libraries were sequenced on an Illumina HiSeq 2500 (SR50) at The Genomics Resource at the Fred Hutchinson Cancer Research Center. Image analysis and base calling were performed using Illumina’s Real Time Analysis v1.18.66.3 software, followed by ‘demultiplexing’ of indexed reads and generation of FASTQ files using Illumina’s bcl2fastq Conversion Software v1.8.4. Reads determined by the RTA software to pass Illumina’s default quality filters were concatenated for further analysis. The FASTQ files were aligned and analyzed using Rockhopper software (McClure et al., 2013). These data have been deposited to the GEO and are accessible using accession number GSE150188.

### qRT-PCR

Bacteria were grown in the same manner as for RNA-seq except 500µM of each aldehyde tested was added to the culture at mid-log (0.4-0.8 OD_600_). The RNA was extracted by the acidified phenol method as listed above and DNase treated and reverse-transcribed using the iScript Reverse Transcription Supermix (Bio-Rad, #1708840). SYBR Green (Thermo Fisher Scientific # K0223) was then used to amplify genes of interest and CT values and relative expression were normalized using CFX Maestro Software (Bio-Rad #12004110).

### Intracellular RNA extraction

RNA extraction from macrophages was performed as previously described (Sigal, Pasechnek and Herskovits, 2016). J774 macrophages were seeded at a density of 2.0×10^7^ cells/dish in three 150 mm dishes in 30mL media and incubated overnight. The next day, overnight *L. monocytogenes* culture grown at 30°C was washed twice with PBS and added to the cells at a MOI of 50. After one hour, the cells were washed twice with PBS and media containing gentamicin was added. Eight hours post-infection, cells were washed once with PBS and lysed by addition of cold nuclease-free water. Lysate was collected by scraping and centrifugation at 800g for 3 min at 4°C. Supernatants were passed through 0.45 µm filters in a vacuum apparatus, and filters were collected in conical tubes. Filters were vortexed with 650µL sterile AE buffer for one minute and centrifuged briefly. Bacteria-containing AE buffer was collected and used for immediate RNA extraction as described above.

### Survival assays Firmicute panel

Bacteria were inoculated overnight in TSB and grown at 37°C. The next day the bacteria were subcultured 1:1000 into fresh TSB and allowed to reach mid-log (0.4-0.8 OD_600_). At mid-log the ODs of the bacteria were normalized to OD1, washed twice in sterile PBS and resuspended in sterile PBS. Then the bacteria were diluted 1:100 into sterile PBS in Eppendorf tubes and various concentrations of 4-HNE were added. The bacteria were then placed at 37°C for an hour. After an hour, the bacteria were plated on LB plates to enumerate CFU.

### *L. monocytogenes* competition experiments

Colonies of *L. monocytogenes* were picked off BHI plates and inoculated into 2mL TSB which was then grown shaking at 37°C to mid-log (0.5-0.8 OD_600_). At mid-log the bacteria were normalized to OD_600_ 1, washed twice in sterile PBS and resuspended in sterile PBS. Then the bacteria were diluted 1:100 into sterile PBS and appropriate strains were mixed together in a 1:1 ratio, after which 640µM of 4-HNE or vehicle (ethanol) was added. The bacteria were then placed at 37°C for one hour. 2x concentrated TSB media was added to the bacterial-PBS solution and the cells recovered for an hour at 37°C. The bacteria were then plated on BHI-streptomycin and BHI-chloramphenicol plates for competitive strain differentiation. CFUs were enumerated after 24-48 hours of growth.

### Purified protein expression and enzyme kinetics

*rha1* and *rha2* ORFs were cloned into pET20B vectors and transformed into BL21 *E. coli* and grown overnight in LB-ampicillin. The overnights were subcultured 1:100 in 2L baffled flasks until mid-log (0.5-0.8 OD_600_) after which 0.5mM IPTG was added. The *pET20b::rha2 E. coli* were induced for 4 hours at 37°C shaking. 0.1% w/v riboflavin was added to *E. coli pET20B::rha1* and the flask was moved to grow at 17°C for 18 hours shaking. Upon completion, the bacteria were spun down, resuspended in buffer A (30mM K2HPO4, 300mM NaCl, pH 8) and sonicated on ice with a large sonicator tip at 80% power 1 sec on 1 sec off for 30 seconds total. They were then spun down at 15,000 g for 45 minutes at 4°C and the supernatant was passed over a nickel resin column (Thermo Fisher Scientific, # PI88222) and eluted using buffer B (30mM K2HPO4, 300mM NaCl, 500mM Imidazole, pH 8). The final protein was then transferred into PBS using a desalting column (Bio-Rad #7322010). For purification of Rha1 10µM FMN (Sigma #F2253) was added at every step of purification.

Enzyme turnover assays were performed at 37°C in 96 well clear bottom plates (Genesee Scientific, #25-104) in a Synergy HTX plate reader in 200µL PBS using 200µM NADPH, 0.2µM enzyme and a range of 4-HNE concentrations from 0 to 0.4mM. Rha1 turnover was performed in the presence of 10µM FMN.

### *B. subtilis* growth curves

*B. subtilis* expressing genes of interest on the pHT01 plasmid (Nguyen, Phan, and Schumann, 2007) were struck on LB-chloramphenicol plates on day 1 and grown overnight at 30C. On day 2, colonies were re-struck on LB chloramphenicol IPTG 1mM LB plates overnight at 30C. On day 3, biomass was scraped and processed as described in the “bacterial culturing” section above. The bacteria were then normalized to an OD_600_ of 1 and inoculated 1:100 into a 96-well plate containing TSB chloramphenicol and 0.5mM IPTG. 4-HNE was then added to the bacteria at various concentrations and the bacteria were allowed to grow at 37°C in Synergy HTX plate reader for 12 hours with shaking.

### *B. subtilis* 4-HNE dot blot

*B. subtilis* were grown and processed in the same manner as for the growth curves above. Upon OD_600_ 1 normalization the bacteria were resuspended in TSB-chloramphenicol with 0.5mM IPTG and 250µL of this mix was transferred to a sterile Eppendorf tube into which 640µM 4-HNE was added. The tubes were then incubated at 37°C for 3 hours. At 3 hours, the bacteria were spun down at 17,000g for 1 minute. The supernatant was then aspirated and the pellet flash frozen in liquid nitrogen. At this point, the bacteria were stored at -80C until further processing. Once removed from the -80C, the bacteria were thawed at room temperature and resuspended in 250µL PBS. The bacteria were then sonicated using a narrow tip sonicator at 20% power, 1sec on 1sec off for 10 seconds and placed on ice. The bacteria were then spun at 4°C at 5,000g for 10 minutes. The subsequent lysate was transferred to fresh Eppendorf tubes containing Halt Proteinase and Phosphatase Inhibitor (Thermo Fisher Scientific, #78442) and stored at -80C until use. For the dot blots, the protein concentration was normalized using BCA (Fisher Scientific, #PI23227) and 3µg in 3µL was spotted onto nitrocellulose membrane (Bio-Rad, #1620115). The nitrocellulose was then dried, blocked for 45 minutes in 5% dry milk, washed three times with TBS-T and primary 4-HNE antibody was added at 1:200 dilution (Abcam, #ab46545). The antibody was left on overnight at 4°C rocking. The primary antibody was then washed off with TBS-T three times and secondary antibody (Licor, #926-32211) was added at 1:8000 for 45 minutes at RT. The secondary was then washed off with TBST twice, TBS once, and then the blot was imaged on a Licor Odyssey Fc. Relative densitometric analysis was performed using Licor Image Studio software.

### *L. monocytogenes* macrophage infection

0.5 × 10^6^ primary murine macrophages from WT C57BL/6 mice were plated in BMM media (DMEM with 10% heat inactivated fetal bovine serum, 1mM L-glutamine, 2mM sodium pyruvate and 10% L929-conditioned medium) in a tissue culture treated 24-well dish (Greiner Bio, #662165) with the addition of 100ng recombinant murine IFN-γ (Peprotech, #315-05) for 18 hours. Inoculants of *L. monocytogenes* were statically grown at 30°C overnight, washed twice with sterile PBS and resuspended in PBS before an MOI of 0.1 was added to each macrophage well. The cells were left to sit for an hour after which all wells were washed twice with PBS and gentamicin was added to all but one well, which was lysed in 500µL water and plated for CFU on LB plates. The remainder of the wells were washed twice with PBS and then lysed and plated for CFU at hours 2, 6 and 9 post-infection.

### *B. subtilis* macrophage survival assay

*B. subtilis* were grown on LB-chloramphenicol 1mM IPTG plates and processed into LB chloramphenicol as described above. Once the bacteria were normalized to OD_600_ 1 in LB chloramphenicol, an MOI of 100 was added to 0.5 × 10^6^ primary murine macrophages from WT C57BL/6 or C57BL/6 deficient for phox (*gp91*^*phox-/-*^) mice (The Jackson Laboratory, stock # 002365) that have been activated using 100ng/well recombinant murine IFN-γ (Peprotech, #315-05) for 18 hours. The cells were spinfected at 200g for 5 minutes. At 1.5 hours, cells were washed 2x with sterile PBS and gentamicin was added to the cells. pBMMs were lysed in 500µL cold water at hours 2, 6, 7, and 8 and plated for CFU. Colonies were enumerated after overnight growth at 30°C on LB plates.

---

**Supplemental Fig 1.**
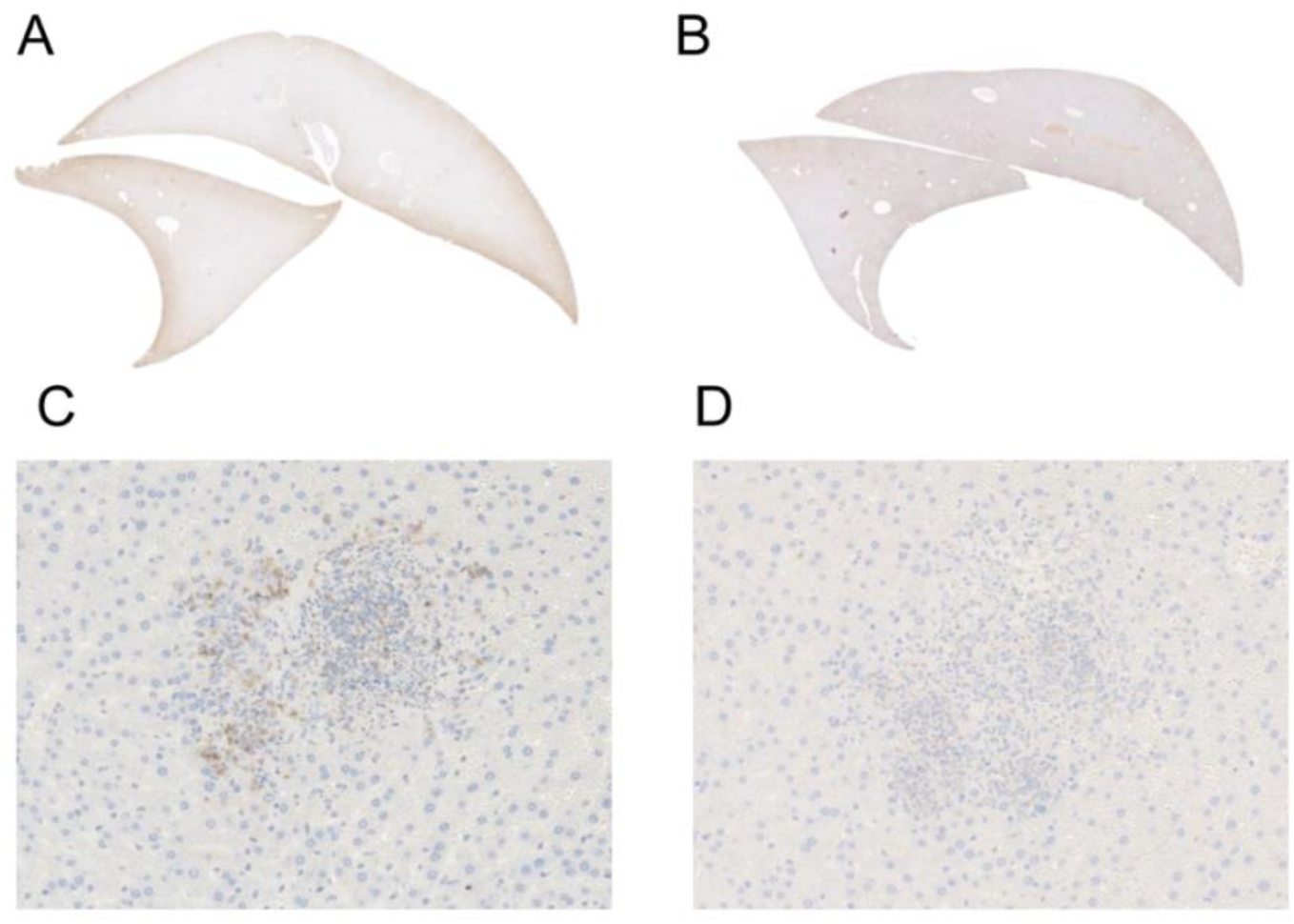
4-HNE accumulation in the liver during 48-hour murine infection by GFP_+_ *L. monocytogenes* assessed by immunohistochemistry analysis. A. Liver (infected) anti-4-HNE B. Liver (uninfected) anti-4-HNE C. Liver (infected) anti-GFP 10X magnification D. Liver (infected) anti-HNE 10X magnification

**Supplemental Fig 2.**
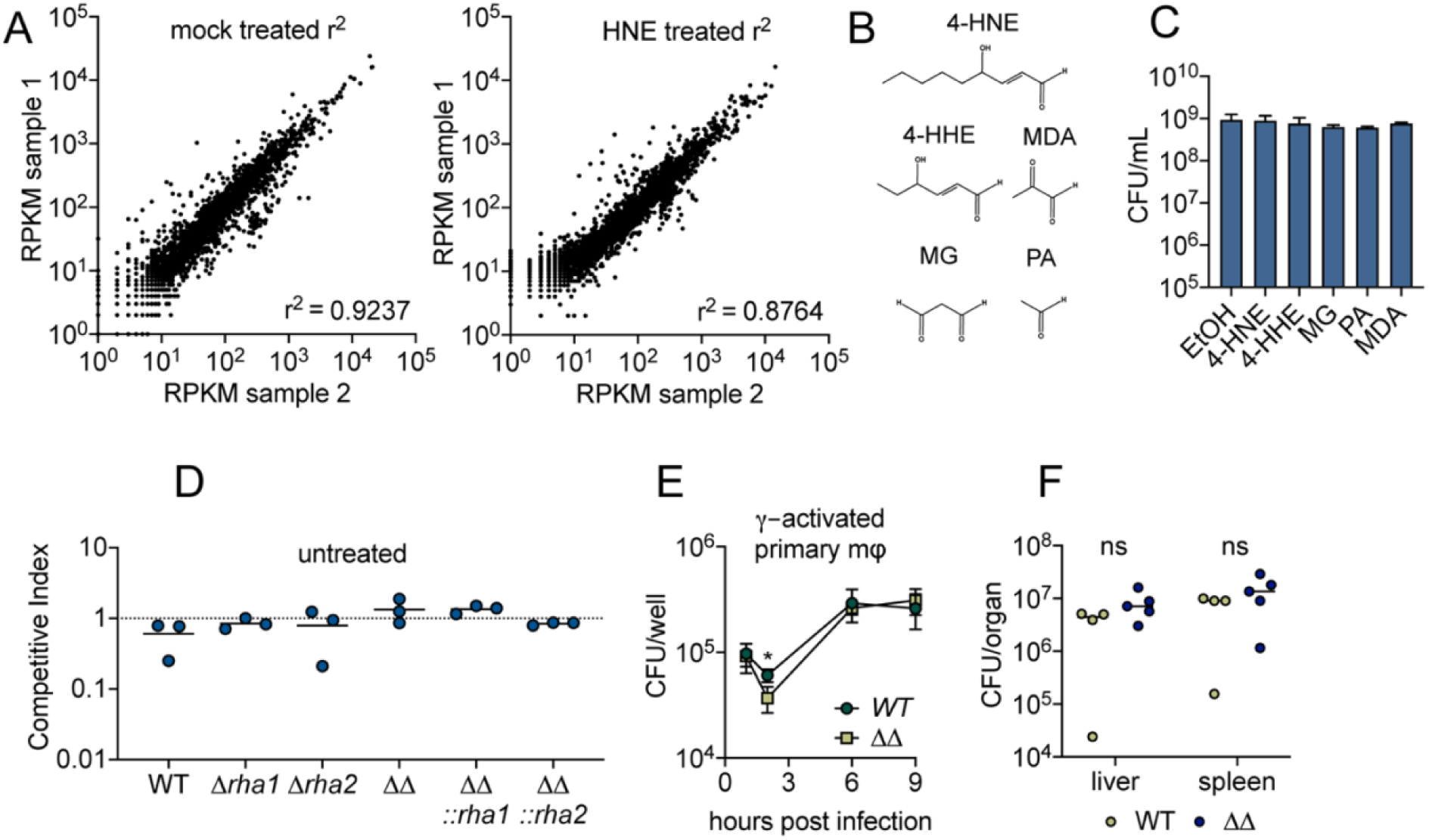
A. Global gene expression of midlog *L. monocytogenes* in TSB treated with ethanol mock control (panel 1) or 640 µM 4-HNE (panel 1) for 20 minutes. RPKM: reads per kilobase million. B. Aldehyde structures used in the experiment. 4-HNE: 4-hydroxy-2-nonenal, 4-HHE: 4-hydroxy-2-hexenal, MDA: malondialdehyde, MG: methylglyoxal, PA: propionaldehyde C. CFU post treatment with sublethal concentrations of listed aldehydes. Performed in technical duplicate. D. Competitive index of 1 hour mock treated (ethanol) *L. monocytogenes* in PBS. E. CFU/well of WT and *Δrha1Δrha2 L. monocytogenes* in 100ng recombinant murine IFN-γ activated WT primary murine macrophages. Performed in biological triplicate. F. CFU/organ of WT and *Δrha1Δrha2 (ΔΔ) L. monocytogenes* at 48 hours intravenous murine infection. Statistics in (E) are unpaired t-tests comparing *WT* and *ΔΔ L. monocytogenes* CFU at hour 2 post infection. Statistics in (F) are unpaired t-tests comparing *WT* and *ΔΔ L. monocytogenes* CFU within each organ. Error bars are mean +/- SD. ns > 0.05; *, p < 0.05. In figure (F), the line is drawn at the median of data.

**Supplemental Fig 3.**
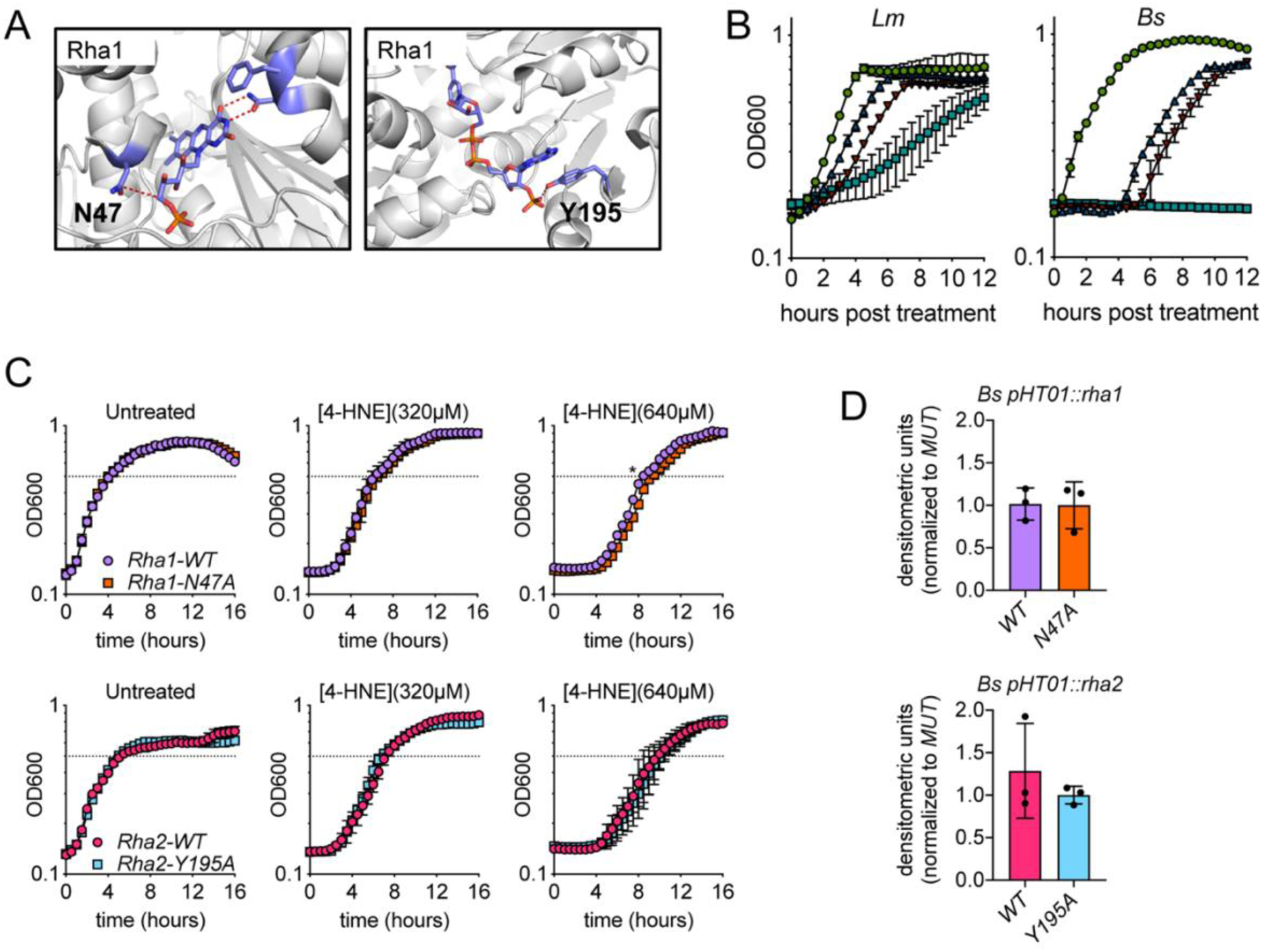
A. Modeling of Rha1 and Rha2 binding pockets with FMN (Rha1) and NADPH (Rha2) with corresponding predicted coordinating amino acid labeled. Coordinating amino acids are labeled. Blue indicates carbon, red indicates oxygen, orange indicates phosphorous. Rha1 is a predicted flavin containing reductase. In comparison between the structure of CLA-ER and the homology model of Rha1 predicted by Phyre2 analysis, several conserved residues lining the flavin binding pocket were identified and the N47A mutant was generated. Rha2 is a predicted NADPH reductase. Similar structural models were generated by Phyre2 analysis and a Y195A mutation in a conserved tyrosine residue predicted to be involved in NADPH co-substrate binding was generated. Figure generated in PyMol. B. *L. monocytogenes* and *B. subtilis* growth curves in TSB after exposure to various concentrations of 4-HNE. (green circles: 0µM, blue triangles: 320µM, red triangles: 640µM, light blue squares: 1280µM). Data performed in technical duplicate and representative of at least three independent experiments. C. Growth curves of *B. subtilis::pHT01* expressing various *rha1* and *rha2* constructs in TSB after exposure to various concentrations of 4-HNE. Dotted line represents OD_600_ 0.5. Performed in biological triplicate. D. 4-HNE dot blots of *B. subtilis::pHT01* expressing various *rha1* and *rha2* constructs after 3 hours of exposure to 640µM 4-HNE in TSB. Performed in biological triplicate. Statistics in (C) is an unpaired t-test comparing the hours to OD 0.5 between *B. subtilis pHT01::rha1-WT* and *pHT01::rha1-N47A*. Error bars are mean +/- SD. *, p < 0.05.

**Supplemental Table 1.**
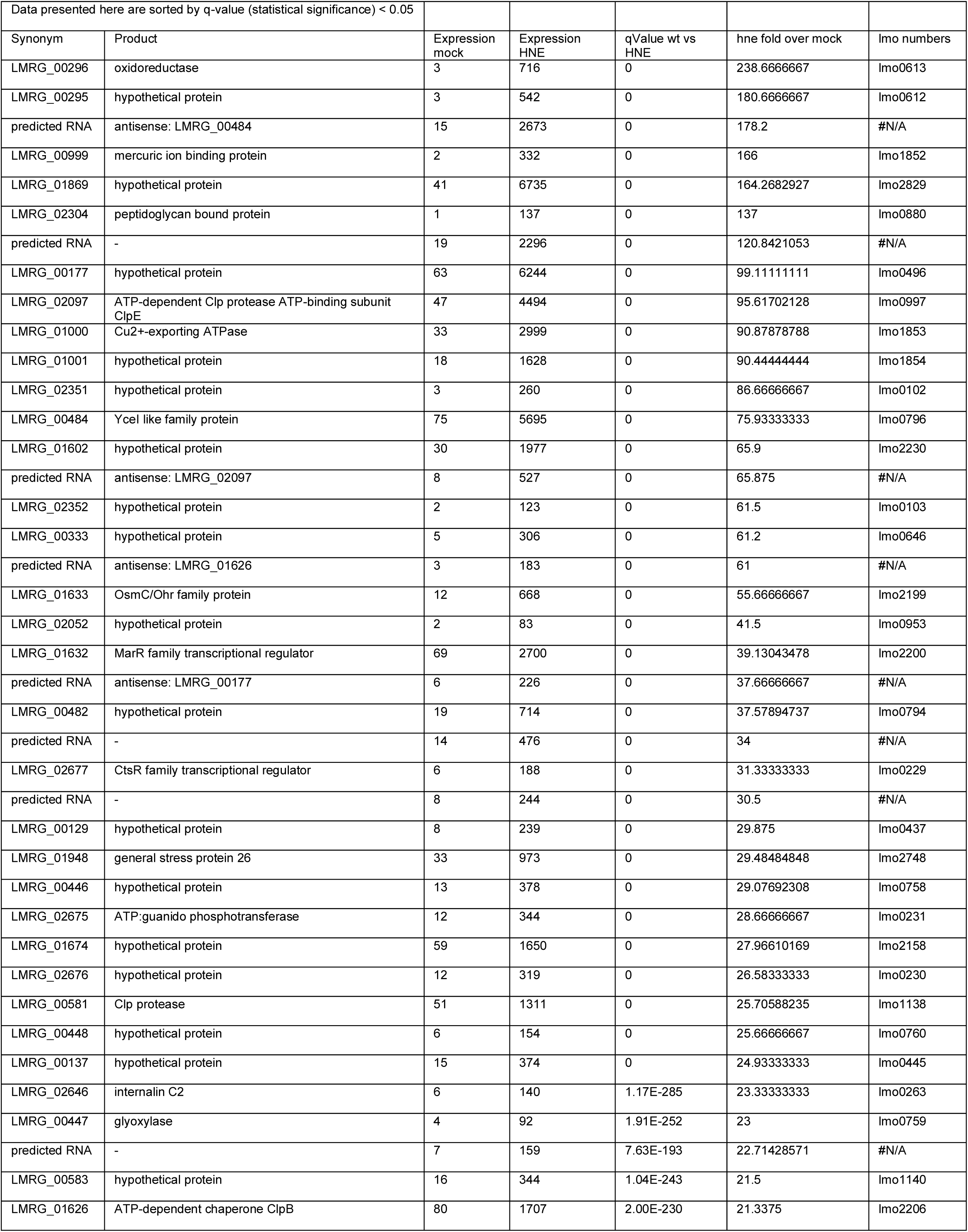
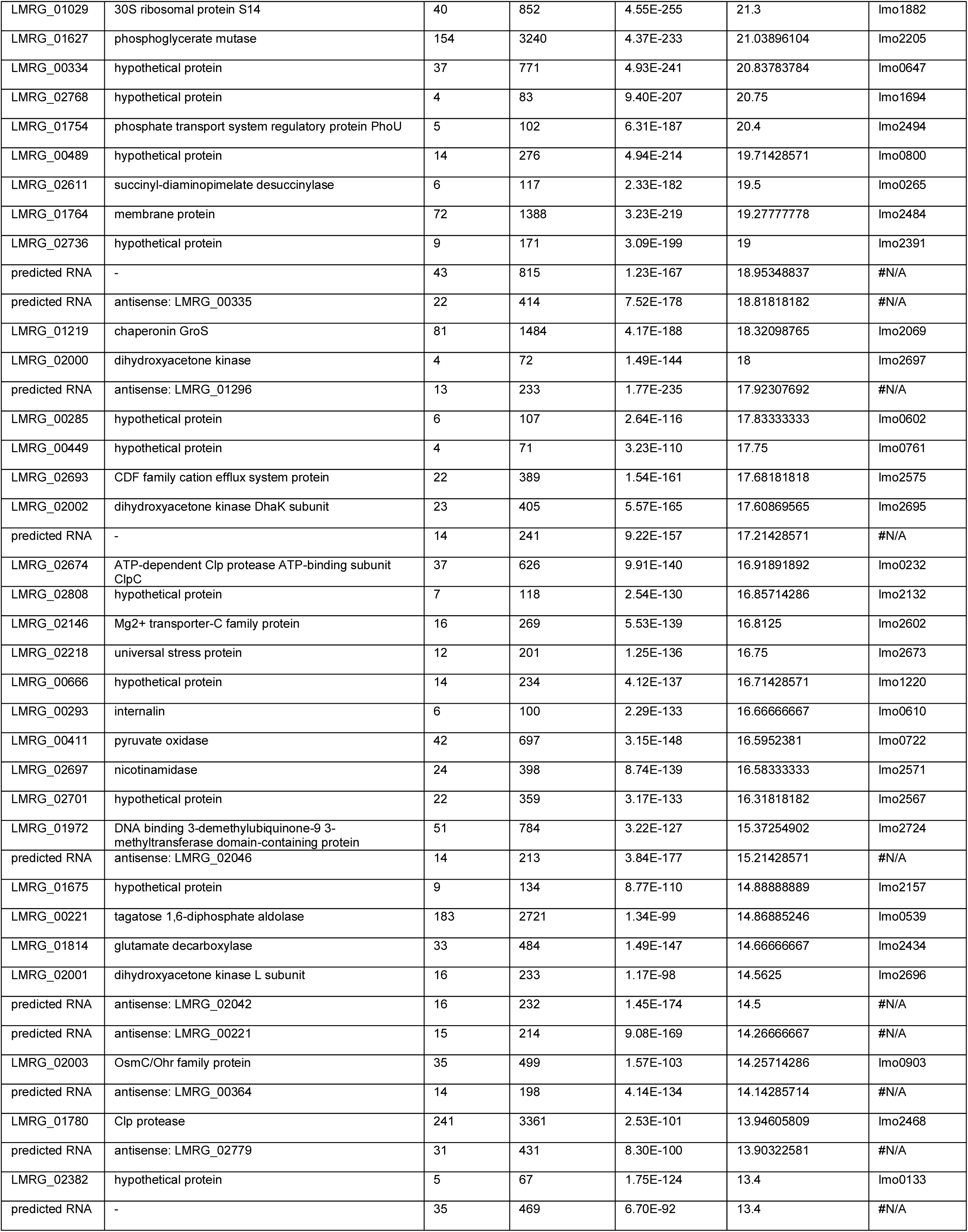
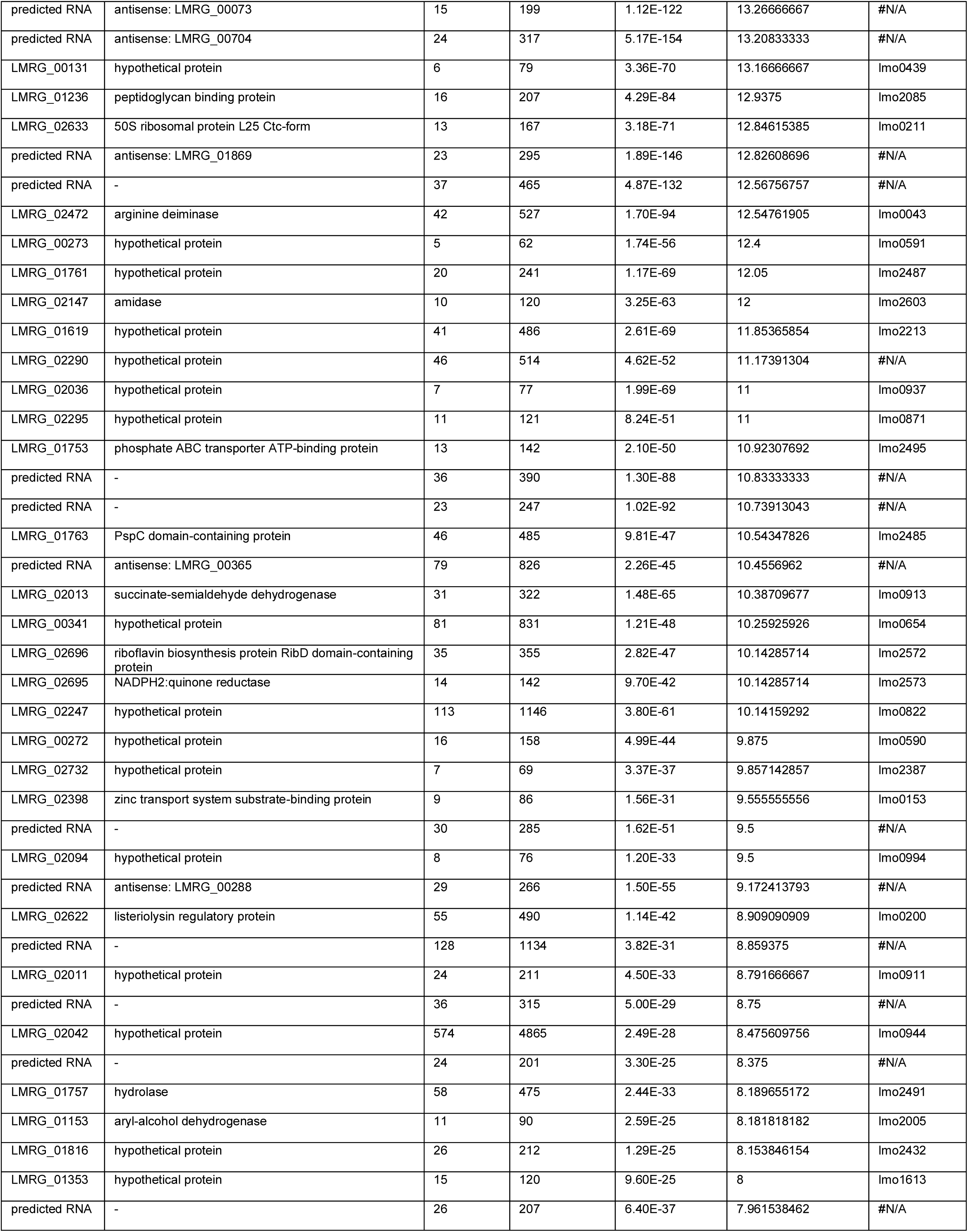
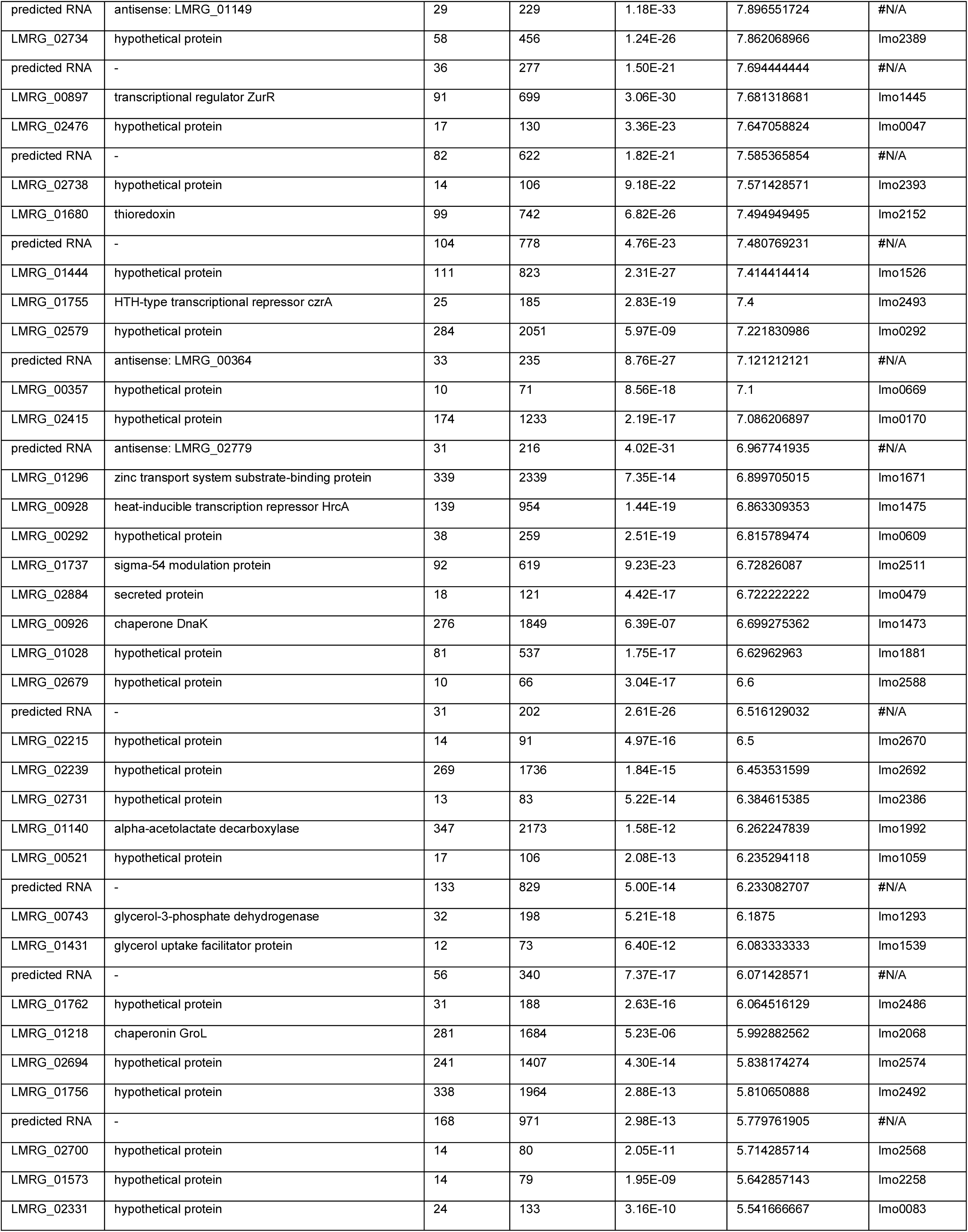
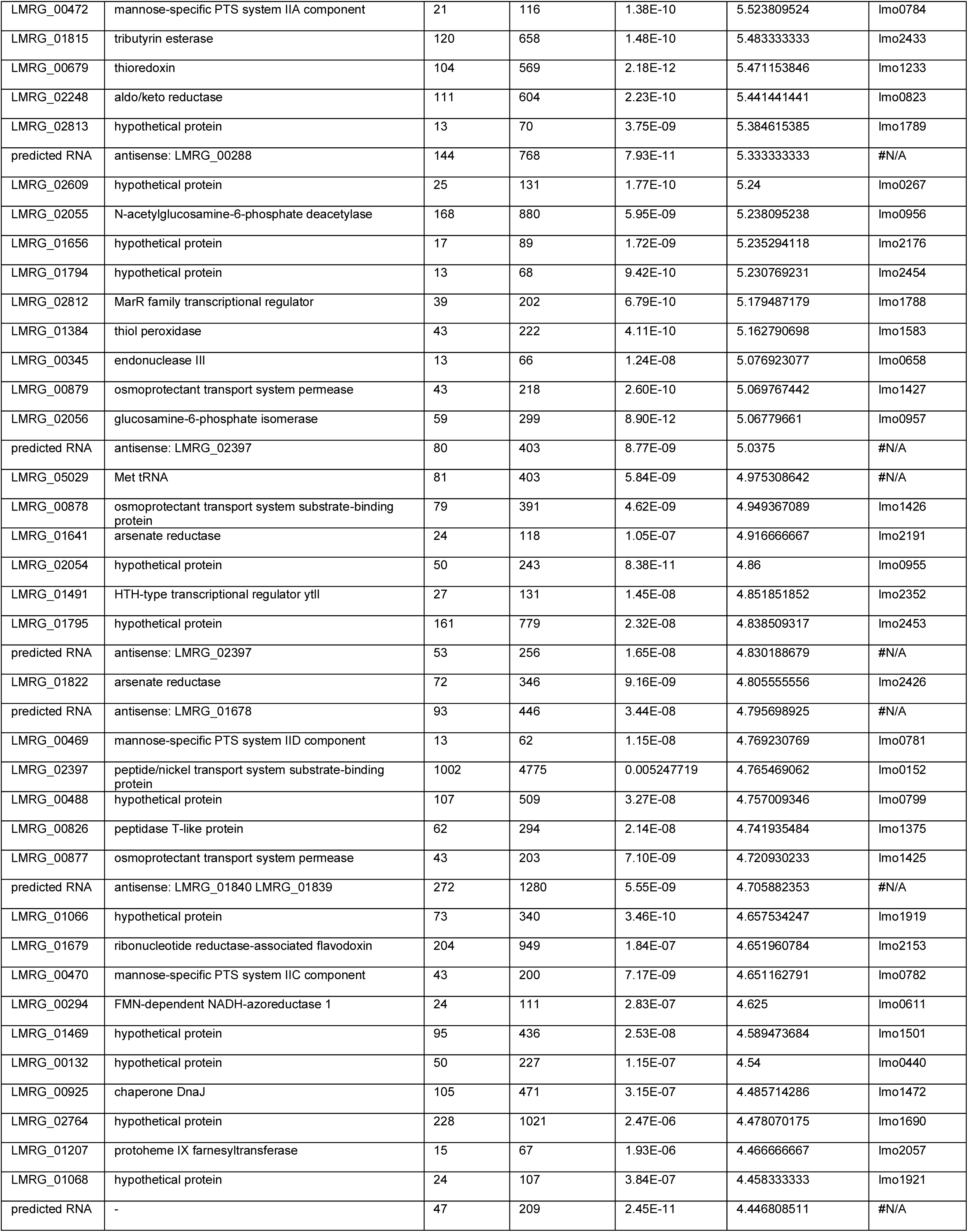
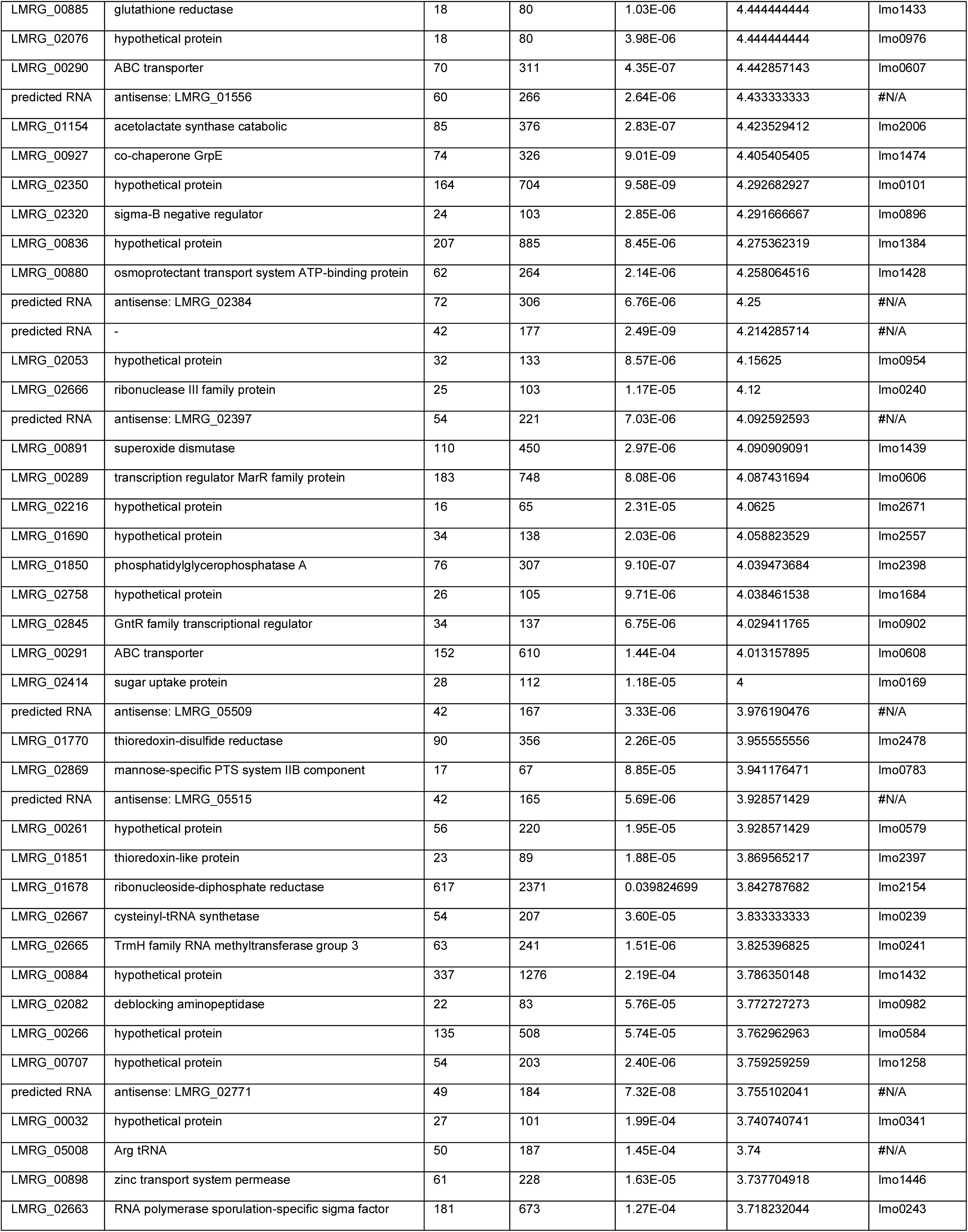
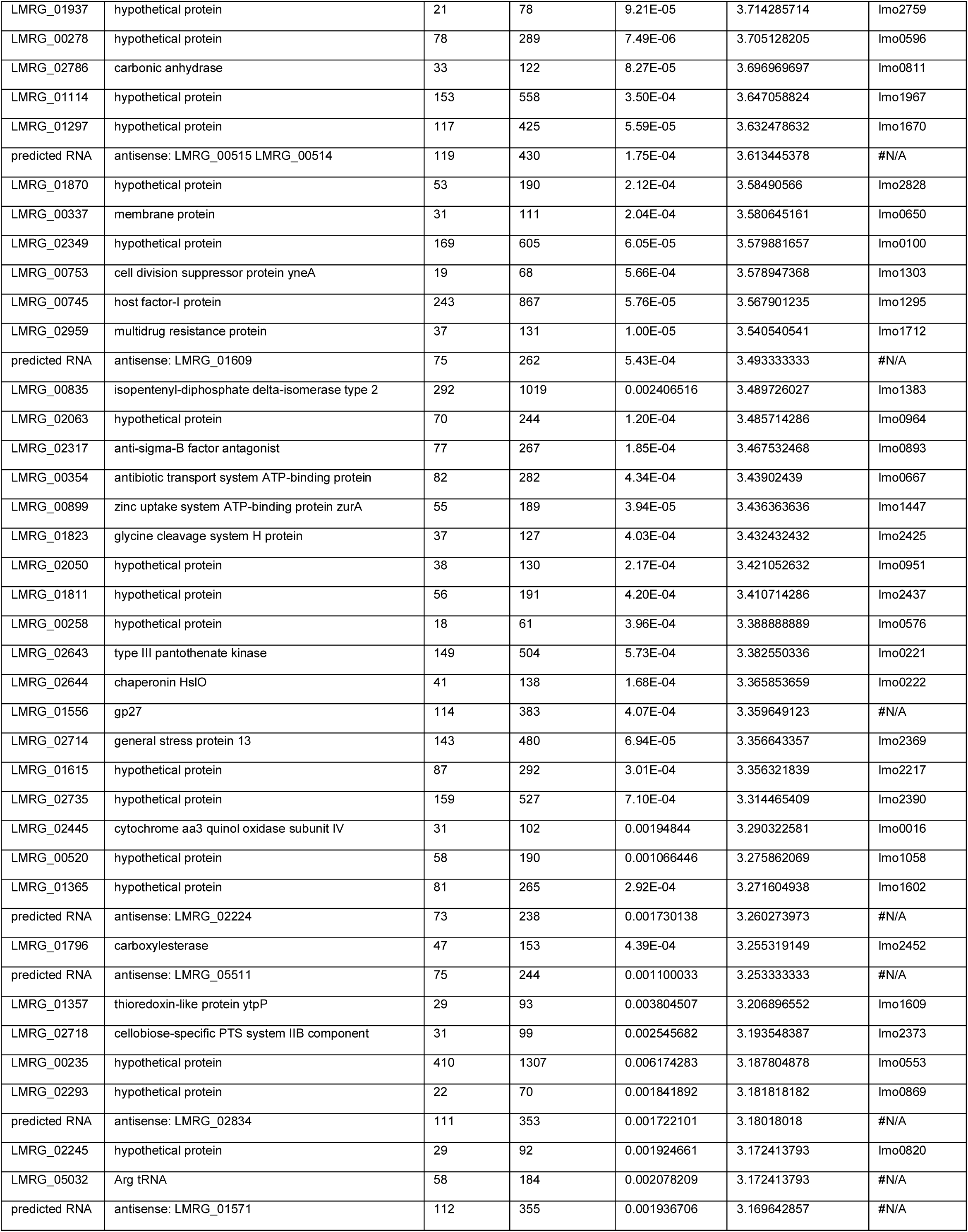
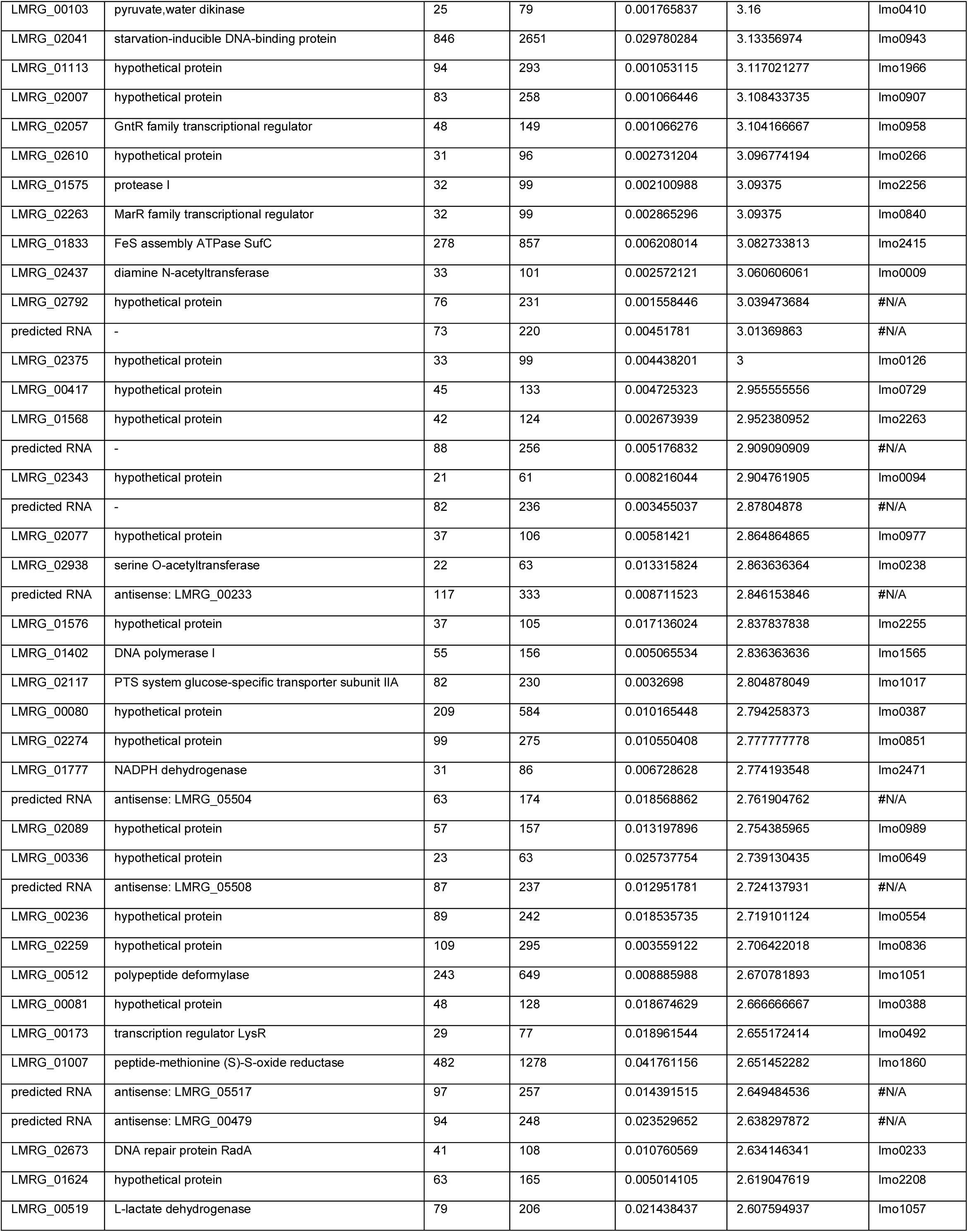
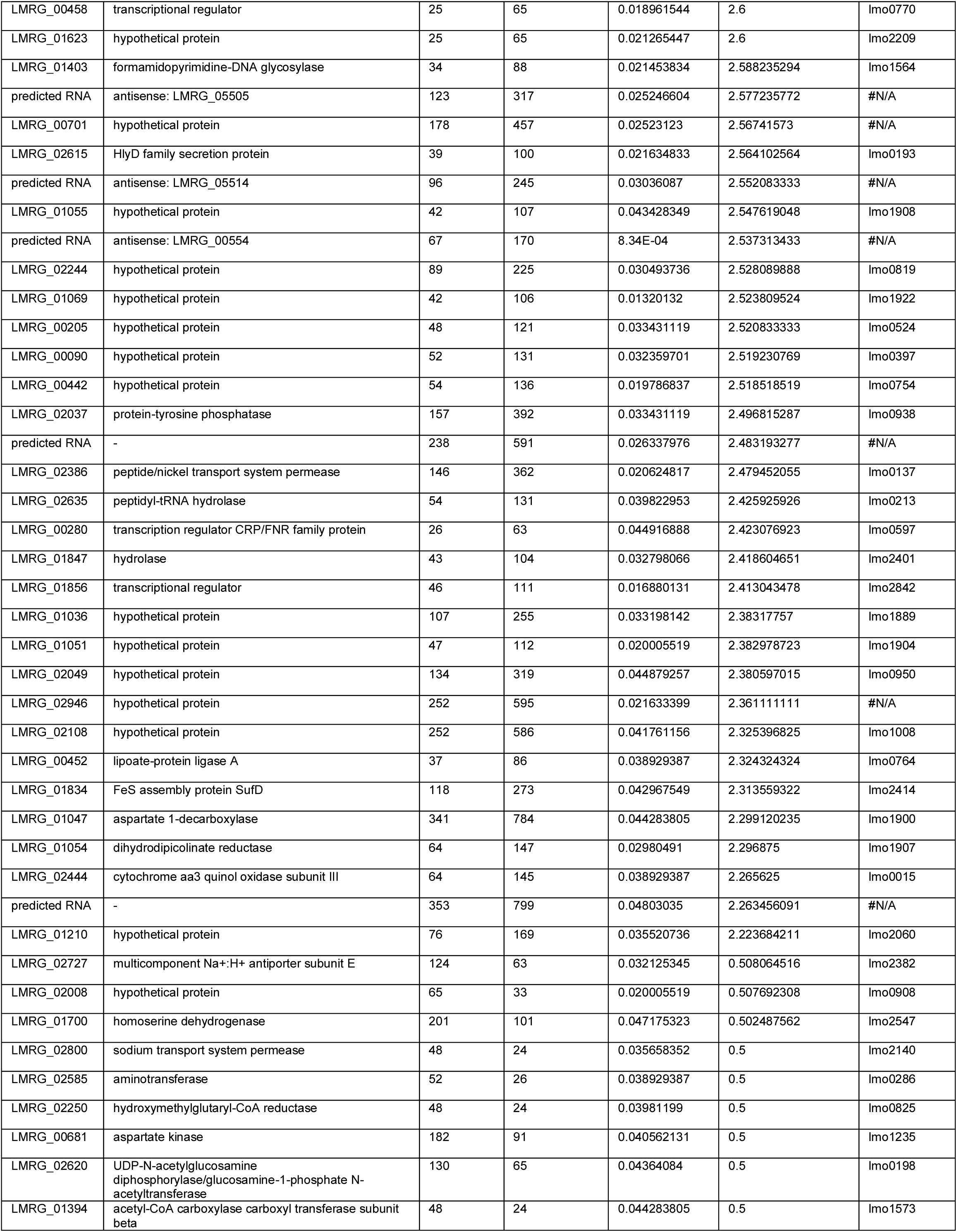
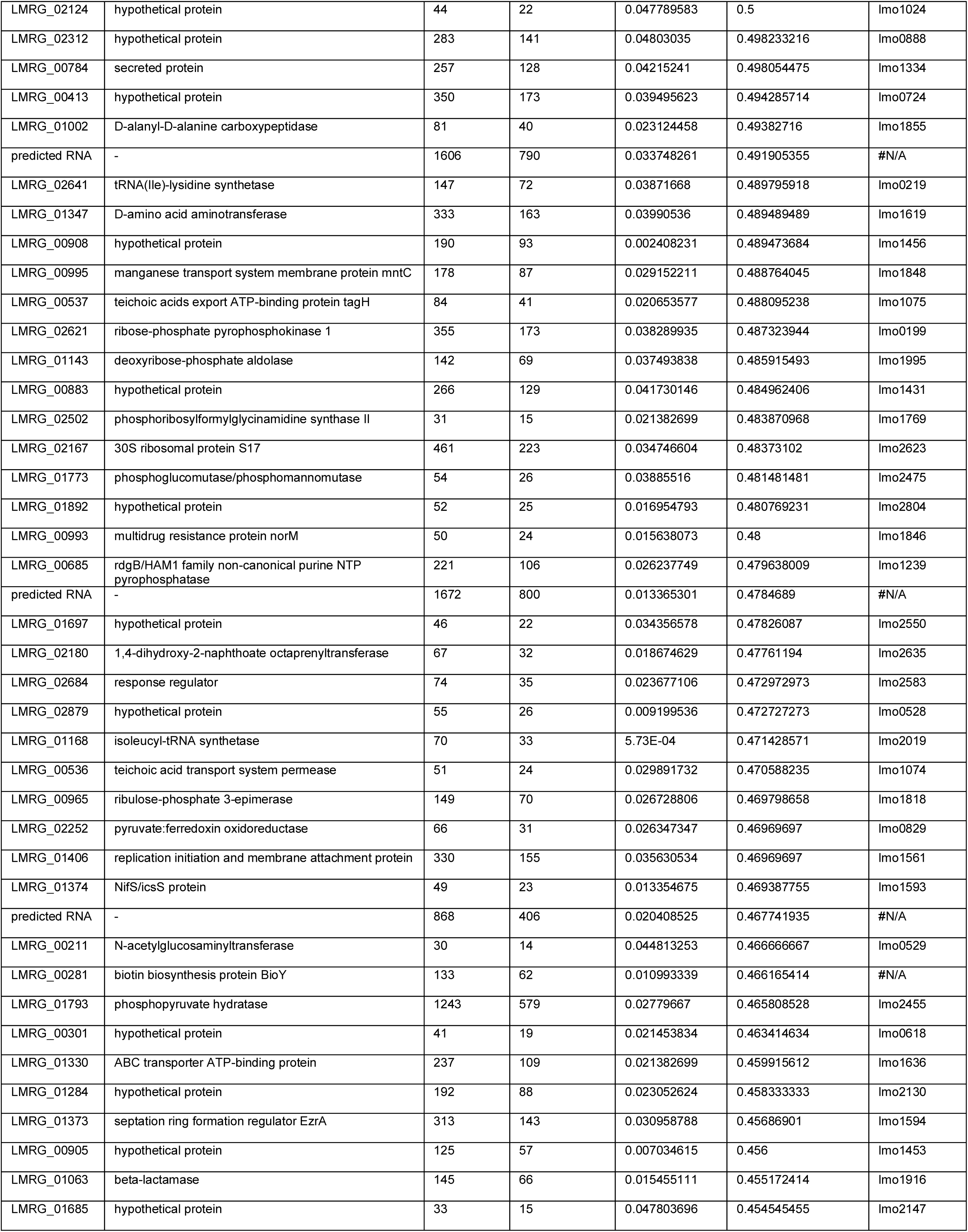
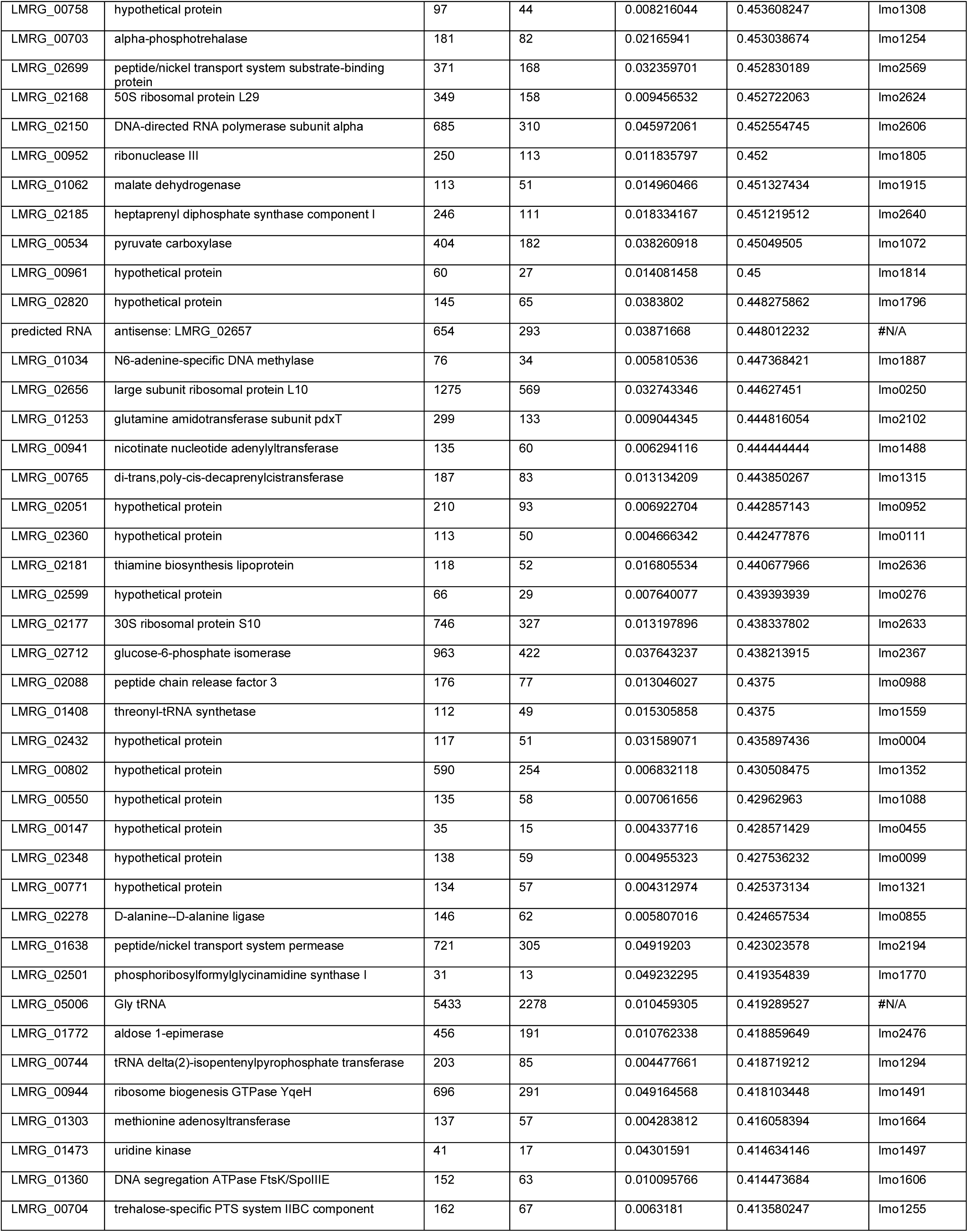
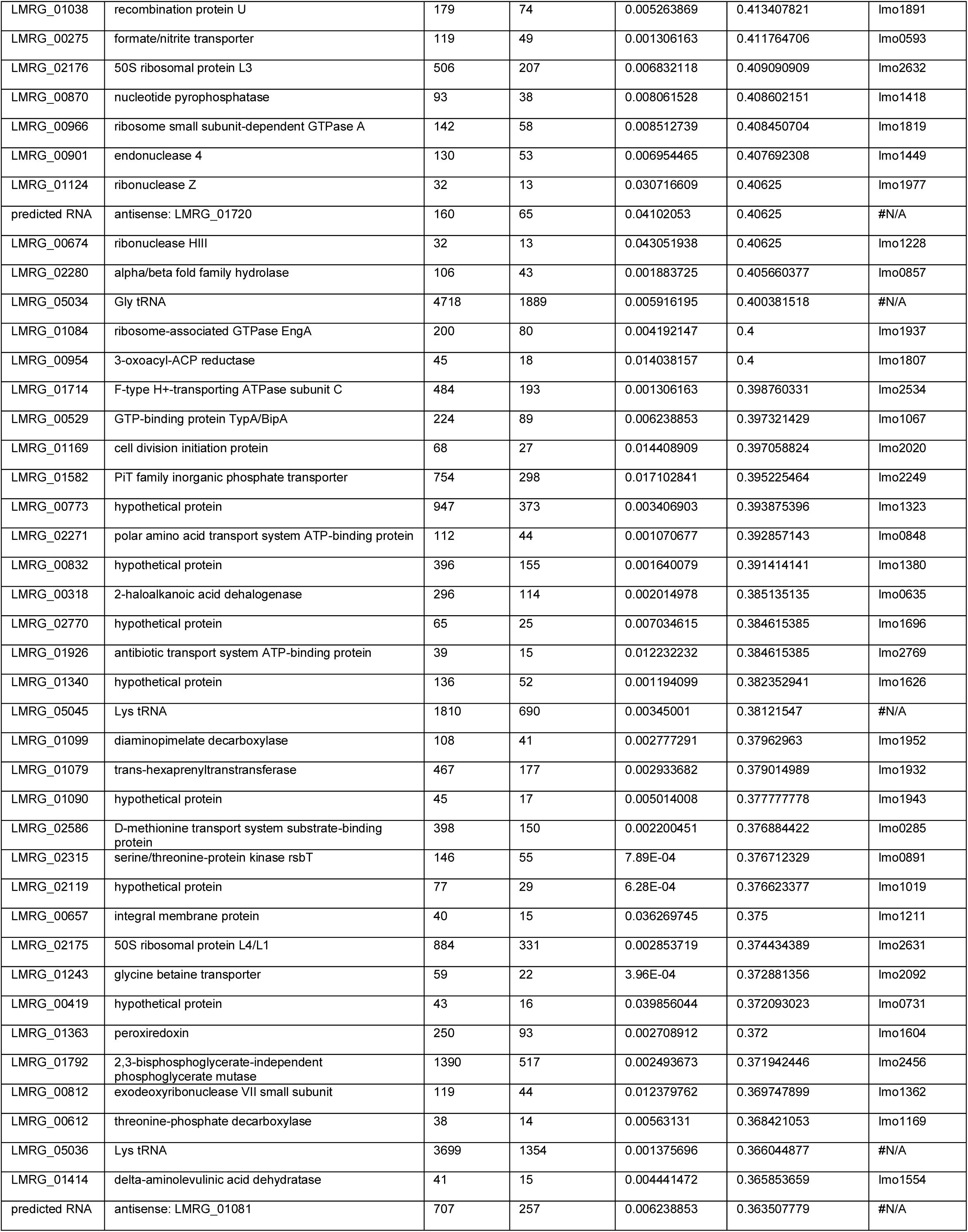
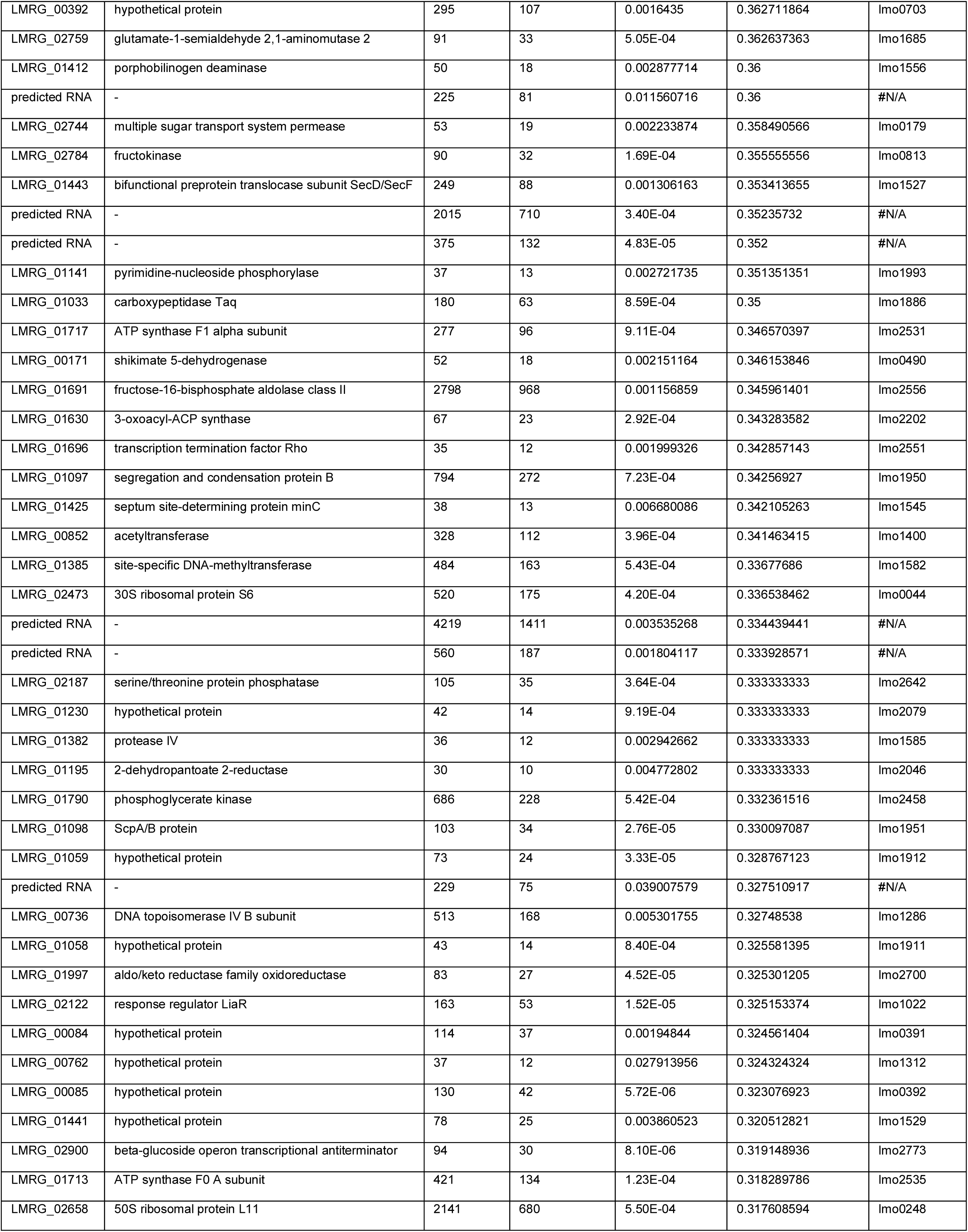
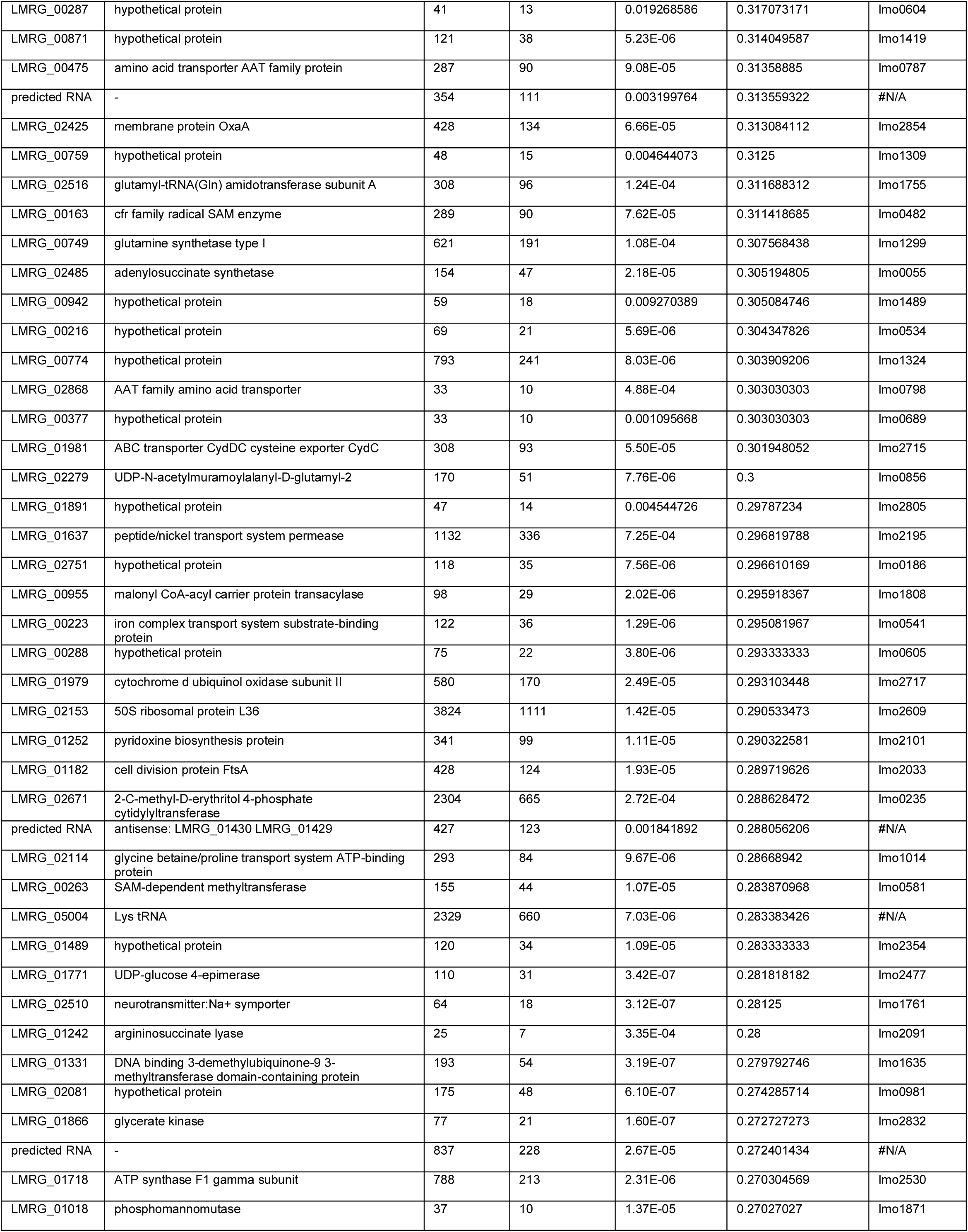
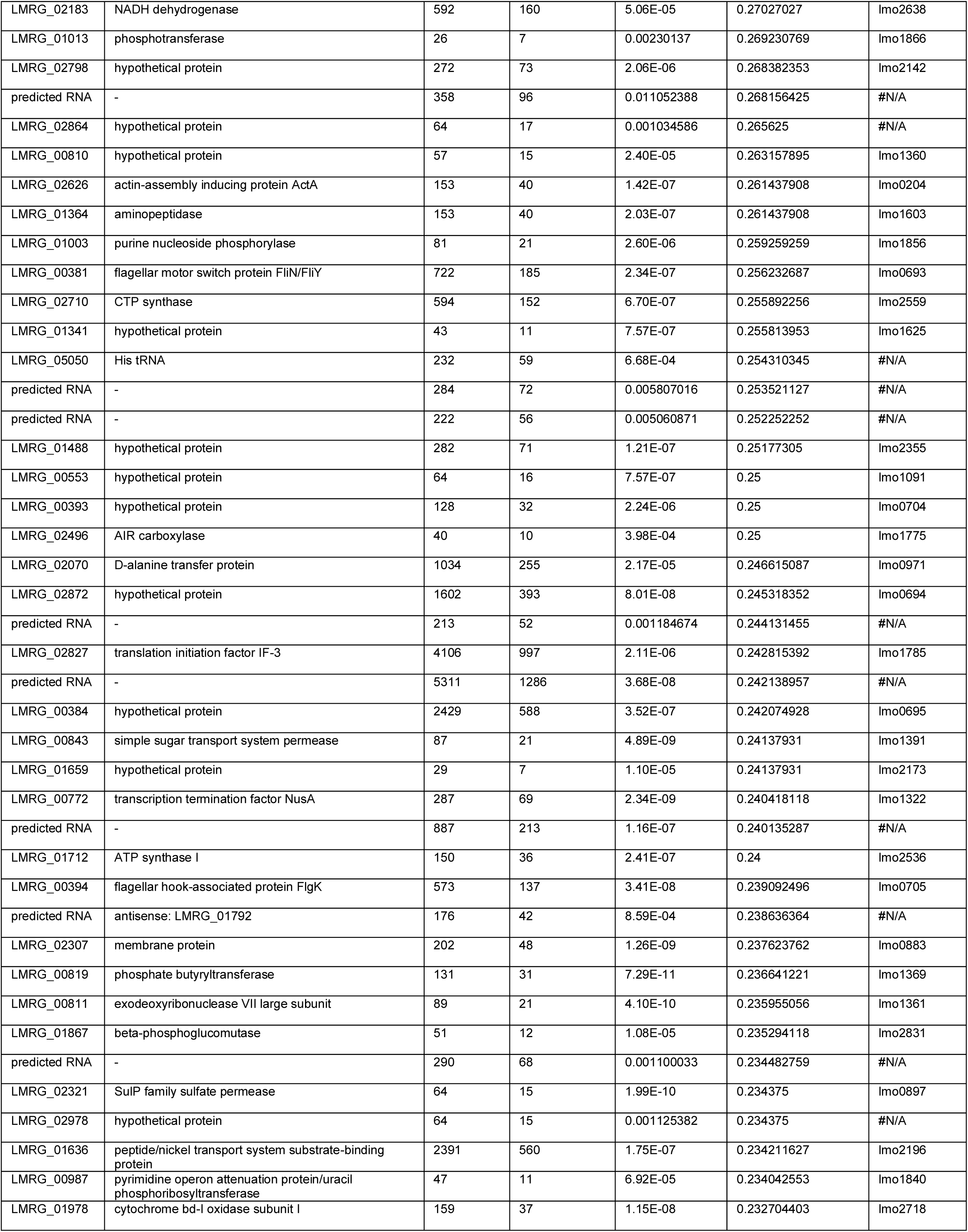
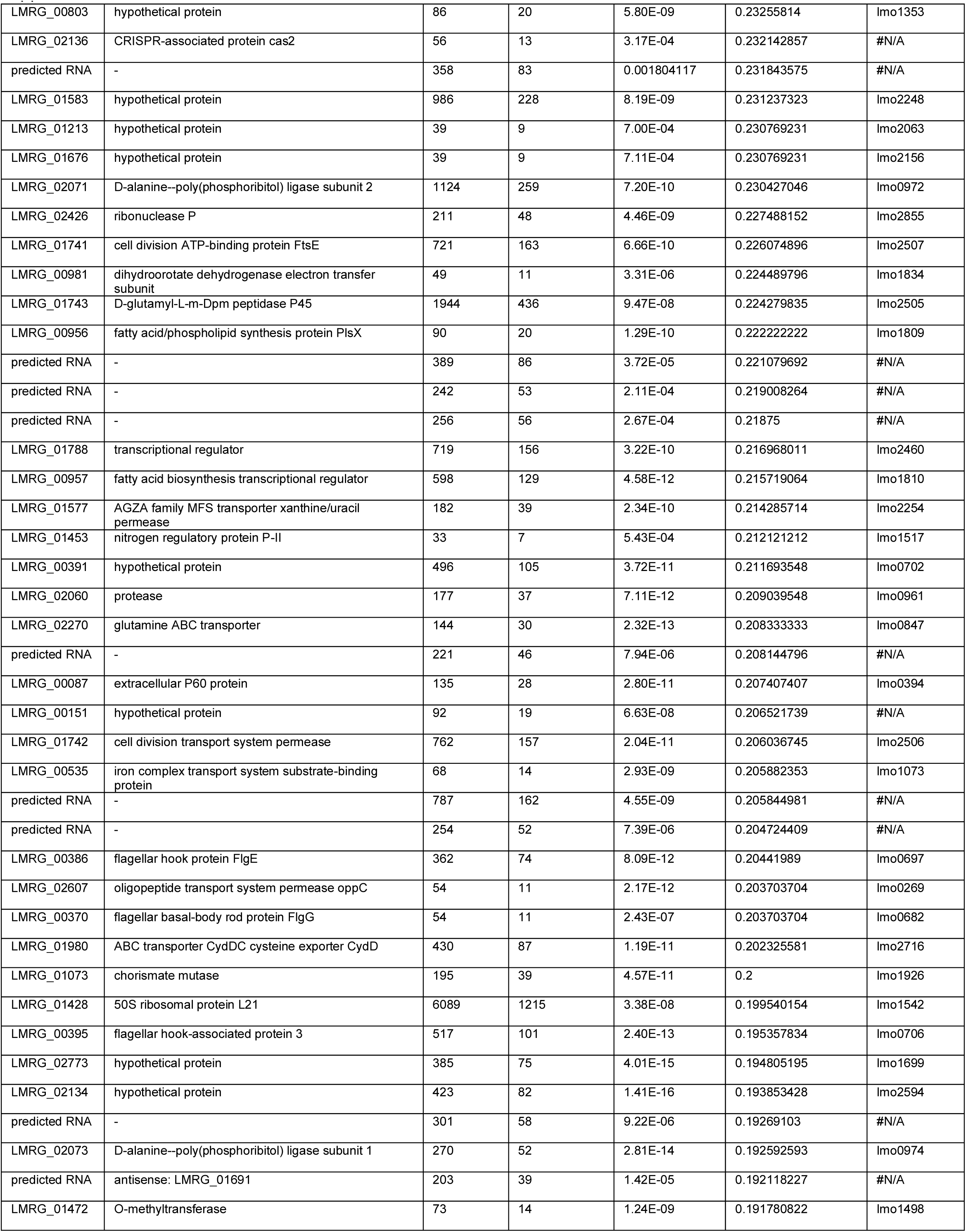
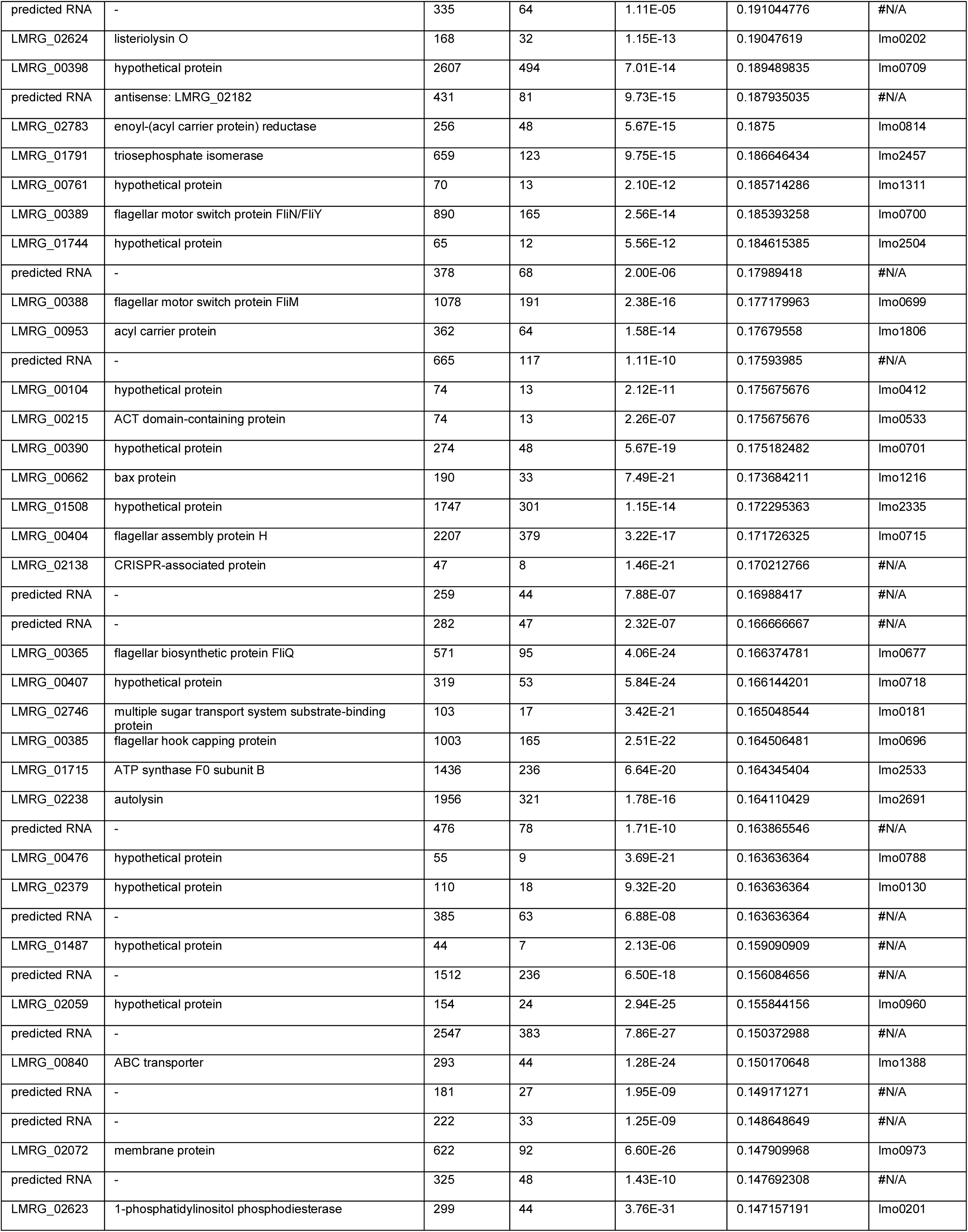
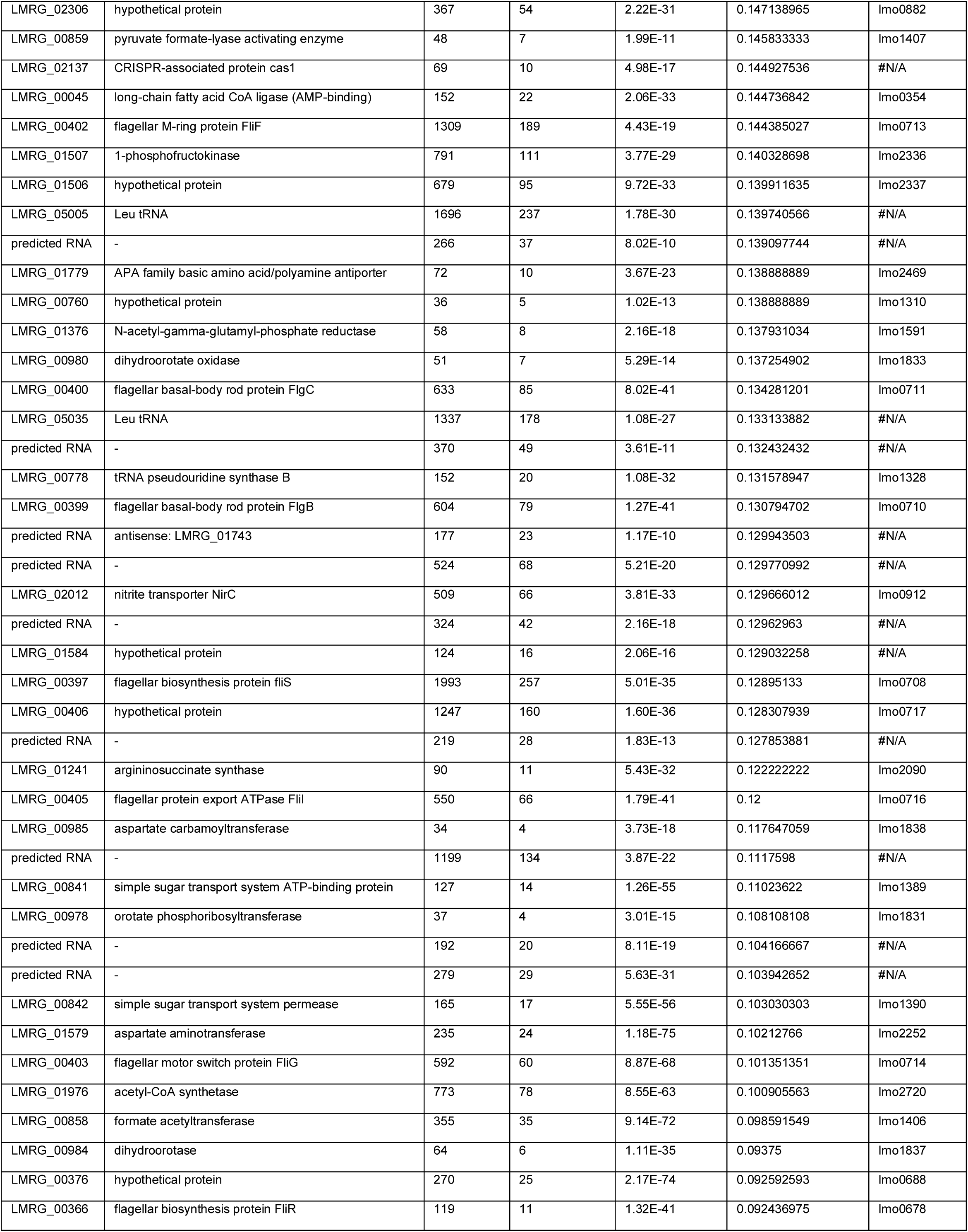
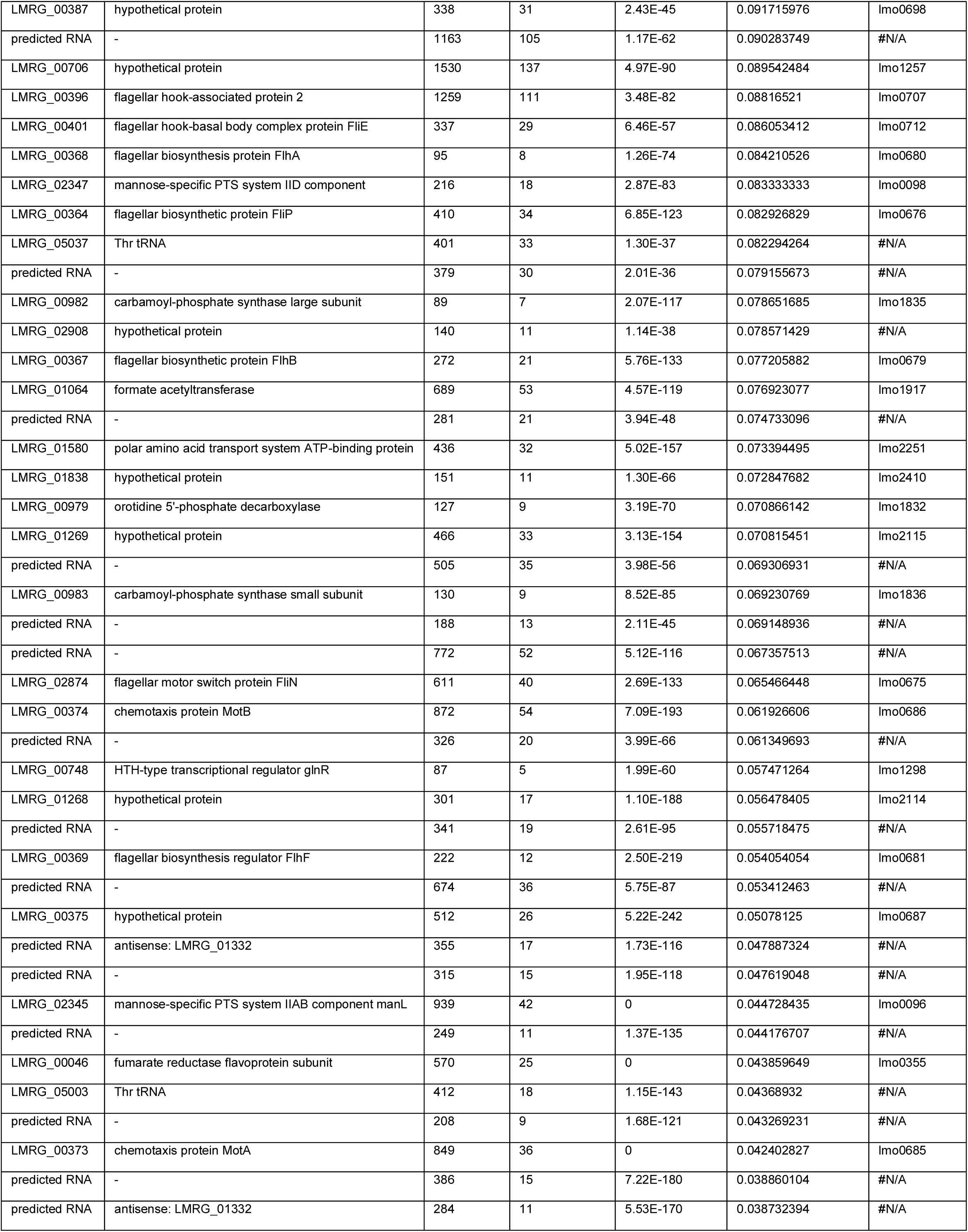
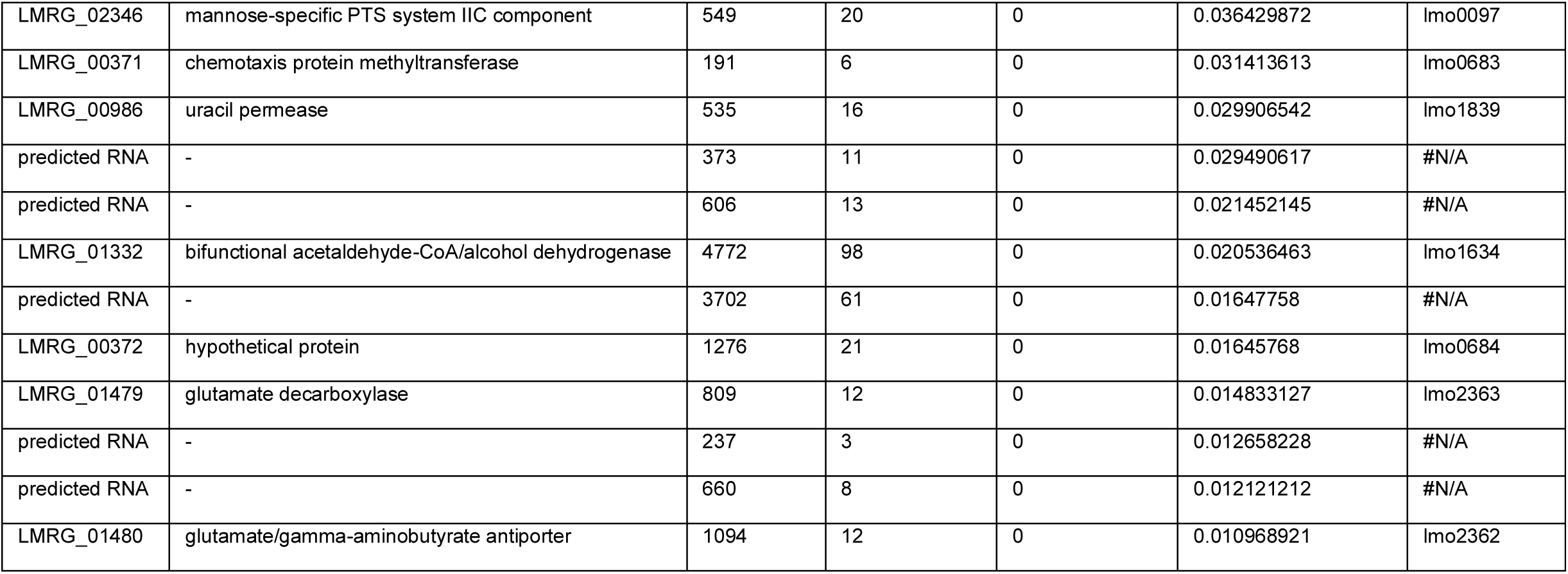

## References

1. Awada, M. et al. (2012) ‘Dietary oxidized n-3 PUFA induce oxidative stress and inflammation: Role of intestinal absorption of 4-HHE and reactivity in intestinal cells’, Journal of Lipid Research. American Society for Biochemistry and Molecular Biology, 53(10), pp. 2069–2080. doi: 10.1194/jlr.M026179.

2. Cybulski, R. J. et al. (2009) ‘Four superoxide dismutases contribute to Bacillus anthracis virulence and provide spores with redundant protection from oxidative stress’, Infection and Immunity. American Society for Microbiology Journals, 77(1), pp. 274–285. doi: 10.1128/IAI.00515-08.

3. Dalleau, S. et al. (2013) ‘Cell death and diseases related to oxidative stress:4-hydroxynonenal (HNE) in the balance’, Cell Death and Differentiation. Nature Publishing Group, pp. 1615–1630. doi: 10.1038/cdd.2013.138.

4. El-Halfawy, O. M. et al. (2017) ‘Antibiotic capture by bacterial lipocalins uncovers an extracellular mechanism of intrinsic antibiotic resistance’, mBio. American Society for Microbiology, 8(2). doi: 10.1128/mBio.00225-17.

5. Esterbauer, H., Schaur, R. J. and Zollner, H. (1991) ‘Chemistry and biochemistry of 4-hydroxynonenal, malonaldehyde and related aldehydes’, Free Radical Biology and Medicine. Pergamon, pp. 81–128. doi: 10.1016/0891-5849(91)90192-6.

6. Fang, F. C. (2004) ‘Antimicrobial reactive oxygen and nitrogen species: Concepts and controversies’, Nature Reviews Microbiology. Nature Publishing Group, pp. 820–832. doi: 10.1038/nrmicro1004.

7. Galley, H. F. and Webster, N. R. (1998) ‘Nitric oxide in a nutshell: Genetics, physiology and pathology’, Current Anaesthesia and Critical Care. Churchill Livingstone, 9(4), pp. 209–213. doi: 10.1016/S0953-7112(98)80057-4.

8. Hanna, V. S. and Hafez, E. A. A. (2018) ‘Synopsis of arachidonic acid metabolism: A review’, Journal of Advanced Research. Elsevier B.V., pp. 23–32. doi: 10.1016/j.jare.2018.03.005.

9. Hébrard, M. et al. (2009) ‘Redundant hydrogen peroxide scavengers contribute to Salmonella virulence and oxidative stress resistance’, Journal of Bacteriology. American Society for Microbiology Journals, 191(14), pp. 4605–4614. doi: 10.1128/JB.00144-09.

10. Hou, F. et al. (2015) ‘Structure and reaction mechanism of a novel enone reductase’, FEBS Journal. Blackwell Publishing Ltd, 282(8), pp. 1526–1537. doi: 10.1111/febs.13239.

11. Iyengar, R., Stuehr, D. J. and Marlettat, M. A. (1987) Macrophage synthesis of nitrite, nitrate, and N-nitrosamines: Precursors and role of the respiratory burst (L-arginine/’5N enrichment/N-nitrosomorpholine), Proc. Nati. Acad. Sci. USA.

12. Jacobson, M. D. (1996) ‘Reactive oxygen species and programmed cell death’, Trends in Biochemical Sciences. Elsevier Ltd, 21(3), pp. 83–86. doi: 10.1016/S0968-0004(96)20008-8.

13. Lopachin, R. M. and Gavin, T. (2014) ‘Molecular mechanisms of aldehyde toxicity: A chemical perspective’, Chemical Research in Toxicology. American Chemical Society, pp. 1081–1091. doi: 10.1021/tx5001046.

14. Majima, H. J., Nakanishi-Ueda, T. and Ozawa, T. (2002) ‘4-hydroxy-2-nonenal (4-HNE) staining by anti-HNE antibody.’, Methods in molecular biology (Clifton, N.J.). Humana Press, 196, pp. 31–34. doi: 10.1385/1-59259-274-0:31.

15. Mcclure, R. et al. (2013) ‘Computational analysis of bacterial RNA-Seq data’. doi: 10.1093/nar/gkt444.

16. Mol, M. et al. (2017) ‘Enzymatic and non-enzymatic detoxification of 4-hydroxynonenal: Methodological aspects and biological consequences’, Free Radical Biology and Medicine. Elsevier Inc., pp. 328–344. doi: 10.1016/j.freeradbiomed.2017.01.036.

17. Nagafuji, T. et al. (1995) ‘Nitric oxide synthase in cerebral ischemia - Possible contribution of nitric oxide synthase activation in brain microvessels to cerebral ischemic injury’, Molecular and Chemical Neuropathology. Humana Press, 26(2), pp. 107–157. doi: 10.1007/BF02815009.

18. Nathan, C. and Cunningham-Bussel, A. (2013) ‘Beyond oxidative stress: An immunologist’s guide to reactive oxygen species’, Nature Reviews Immunology. Nature Publishing Group, pp. 349–361. doi: 10.1038/nri3423.

19. Nathan, C. F. and Hibbs, J. B. (1991) ‘Role of nitric oxide synthesis in macrophage antimicrobial activity’, Current Opinion in Immunology. Elsevier Current Trends, 3(1), pp. 65–70. doi: 10.1016/0952-7915(91)90079-G.

20. Nguyen, H. D., Phan, T. T. P. and Schumann, W. (2007) ‘Expression vectors for the rapid purification of recombinant proteins in Bacillus subtilis’, Current Microbiology. Springer, 55(2), pp. 89–93. doi: 10.1007/s00284-006-0419-5.

21. Ole Leichert, L. I., Scharf, C. and Hecker, M. (2003) ‘Global characterization of disulfide stress in Bacillus subtilis’, Journal of Bacteriology, 185(6), pp. 1967–1975. doi: 10.1128/JB.185.6.1967-1975.2003.

22. Paradis, V. et al. (1997) ‘In situ detection of lipid peroxidation by-products in chronic liver diseases’, Hepatology. John Wiley and Sons Inc., 26(1), pp. 135–142. doi: 10.1002/hep.510260118.

23. Parsell, D. A. and Lindquist, S. (1993) THE FUNCTION OF HEAT-SHOCK PROTEINS IN STRESS TOLERANCE: DEGRADATION AND REACTIV ATION OF DAMAGED PROTEINS.

24. Patel, R. P. et al. (1999) ‘Biological aspects of reactive nitrogen species’, Biochimica et Biophysica Acta - Bioenergetics. Elsevier, pp. 385–400. doi: 10.1016/S0005-2728(99)00028-6.

25. Pillon, N. J. et al. (2012) ‘The Lipid Peroxidation By-Product 4-Hydroxy-2-Nonenal (4-HNE) Induces Insulin Resistance in Skeletal Muscle through Both Carbonyl and Oxidative Stress’, Endocrinology. Oxford Academic, 153(5), pp. 2099–2111. doi: 10.1210/en.2011-1957.

26. Prost, I. et al. (2005) ‘Evaluation of the Antimicrobial Activities of Plant Oxylipins Supports Their Involvement in Defense against Pathogens 1[W]’, Plant Physiology, 139, pp. 1902–1913. doi: 10.1104/pp.105.066274.

27. Rahman, I. et al. (2002) ‘4-Hydroxy-2-nonenal, a specific lipid peroxidation product, is elevated in lungs of patients with chronic obstructive pulmonary disease’, American Journal of Respiratory and Critical Care Medicine, 166(4), pp. 490–495. doi: 10.1164/rccm.2110101.

28. Roncarati, D. and Scarlato, V. (2017) ‘Regulation of heat-shock genes in bacteria: from signal sensing to gene expression output’, FEMS Microbiology Reviews, 015, pp. 549–574. doi: 10.1093/femsre/fux015.

29. Sayre, L. M. et al. (2002) ‘4-Hydroxynonenal-Derived Advanced Lipid Peroxidation End Products Are Increased in Alzheimer’s Disease’, Journal of Neurochemistry. Wiley-Blackwell, 68(5), pp. 2092–2097. doi: 10.1046/j.1471-4159.1997.68052092.x.

30. Sigal, N., Pasechnek, A. and Herskovits, A. A. (2016) ‘RNA purification from intracellularly grown listeria monocytogenes in macrophage cells’, Journal of Visualized Experiments. Journal of Visualized Experiments, 2016(112). doi: 10.3791/54044.

31. Staerck, C. et al. (2017) ‘Microbial antioxidant defense enzymes’, Microbial Pathogenesis. Academic Press, pp. 56–65. doi: 10.1016/j.micpath.2017.06.015.

32. Stuehr, D. J. and Marletta, M. A. (1985) ‘Mammalian nitrate biosynthesis: Mouse macrophages produce nitrite and nitrate in response to Escherichia coli lipopolysaccharide’, Proceedings of the National Academy of Sciences of the United States of America, 82(22), pp. 7738–7742. doi: 10.1073/pnas.82.22.7738.

33. Uchida, K. (2003) ‘4-Hydroxy-2-nonenal: A product and mediator of oxidative stress’, Progress in Lipid Research. Elsevier Ltd, pp. 318–343. doi: 10.1016/S0163-7827(03)00014-6.

34. Uchida, K. et al. (1994) ‘Michael Addition-Type 4-Hydroxy-2-nonenal Adducts in Modified Low-Density Lipoproteins: Markers for Atherosclerosis’, Biochemistry. American Chemical Society, 33(41), pp. 12487–12494. doi: 10.1021/bi00207a016.

35. Vaughn, S. F. and Gardner, H. W. (1993) ‘Lipoxygenase-derived aldehydes inhibit fungi pathogenic on soybean’, Journal of Chemical Ecology. Kluwer Academic Publishers-Plenum Publishers, 19(10), pp. 2337–2345. doi: 10.1007/BF00979668.

36. Yura, T., Nagai, H. and Mori, H. (1993) ‘Regulation of the Heat-Shock Response in Bacteria’, Annual Review of Microbiology. Annual Reviews, 47(1), pp. 321–350. doi: 10.1146/annurev.mi.47.100193.001541.

37. Zhang, H. and Forman, H. J. (2017) ‘Signaling by 4-hydroxy-2-nonenal: Exposure protocols, target selectivity and degradation’, Archives of Biochemistry and Biophysics. Academic Press Inc., 617, pp. 145–154. doi: 10.1016/j.abb.2016.11.003.

38. Zheng, R. et al. (2014) ‘Differential metabolism of 4-hydroxynonenal in liver, lung and brain of mice and rats’, Toxicology and Applied Pharmacology. Academic Press Inc., 279(1), pp. 43–52. doi: 10.1016/j.taap.2014.04.026.

39. Zimniak, P. (2011) ‘Relationship of electrophilic stress to aging.’, Free radical biology & medicine. NIH Public Access, 51(6), pp. 1087–105. doi: 10.1016/j.freeradbiomed.2011.05.039.

